# Spatiotemporal regulation of the hepatocyte growth factor receptor MET activity by sorting nexins 1/2 in HCT116 colorectal cancer cells

**DOI:** 10.1101/2024.02.10.579658

**Authors:** Laiyen Garcia Delgado, Amélie Derome, Samantha Longpré, Marilyne Giroux-Dansereau, Ghenwa Basbous, Christine L. Lavoie, Caroline Saucier, Jean-Bernard Denault

## Abstract

Cumulative research findings support the idea that endocytic trafficking is crucial in regulating receptor signaling and associated diseases. Specifically, strong evidence points to the involvement of sorting nexins (SNXs), particularly SNX1 and SNX2, in the signaling and trafficking of the receptor tyrosine kinase (RTK) MET in colorectal cancer (CRC). Activation of hepatocyte growth factor (HGF) receptor MET is a key driver of CRC progression. In this study, we utilized human HCT116 CRC cells with *SNX1* and *SNX2* genes knocked out to demonstrate that their absence leads to a delay in MET entering early endosomes. This delay results in increased phosphorylation of both MET and AKT upon HGF stimulation, while ERK1/2 (extracellular signal-regulated kinases 1 and 2) phosphorylation remains unaffected. Despite these changes, HGF-induced cell proliferation, scattering, and migration remain similar between the parental and the *SNX1/2* knockout cells. However, in the absence of SNX1 and SNX2, these cells exhibit increased resistance to TRAIL-induced apoptosis. This research underscores the intricate relationship between intracellular trafficking, receptor signaling, and cellular responses and demonstrates for the first time that the modulation of MET trafficking by SNX1 and SNX2 is critical for receptor signaling that may exacerbate the disease.

## INTRODUCTION

The ability of intracellular trafficking to shape receptor signaling through the successive recruitment of partner proteins *en route* to their destination, like recycling to the plasma membrane or degradation in lysosomes, has been intensely studied [1–3]. This scrutiny reflects the growing understanding that the spatiotemporal control of signaling modulates the outcomes. Receptor tyrosine kinases (RTKs) assemble signalosomes at the plasma membrane but require internalization to fully exert their activity [1,4]. It is now evident that the endosomal system also serves as a pivotal site for signal transduction. Activated RTKs continue to transmit signal from endosomes, yielding responses different from those initiated at the plasma membrane, thereby eliciting distinct physiological responses [5,6]. Notably, this endosomal signaling extends beyond RTKs and has also been shown for G-protein coupled receptors [7,8]. Therefore, studying the intracellular routing of receptors in diseases and their dysregulation in pathologies like cancers may provide valuable clues to identify new therapeutic targets.

Colorectal cancer (CRC) is the second leading cause of cancer-associated death worldwide [9]. An important contributing factor to CRC progression is the aberrant signaling by RTKs, such as MET (c-MET, hepatocyte growth factor/scatter factor receptor). The activation of MET is linked to biological responses that include epithelial growth, morphogenesis, and migration, which becomes deleterious during tumorigenesis and metastasis [10]. Indeed, enhanced expression of MET and its ligand HGF (hepatocyte growth factor) is frequent in metastatic CRC and is generally associated with a poor prognosis [11]. Activation of MET results in the phosphorylation of its C-terminal residues Tyr1349 and 1356, creating a multifunctional docking site for the scaffold proteins GRB2 (growth factor receptor-bound protein 2), SHC (SH2-containing collagen-related proteins), and GAB1 (GRB2-associated binding protein 1). These scaffold proteins mediate the activation of the mitogenic Ras/MAPK (rat sarcoma virus/mitogen-activated protein kinase) and survival PI3K/AKT pathways (phosphatidylinositol 3-kinase/protein kinase B), among others [12]. In addition, the transcription factor STAT3 (signal transducer and activator of transcription 3) interacts with the phosphorylated Tyr1356 of MET [13], promoting epithelial tubulogenesis and anchorage-independent growth [12].

In normal cells, MET signaling is tightly regulated. Upon activation, the receptor is promptly internalized in clathrin-coated vesicles that fuse with early endosomes from where MET is sorted toward either degradation or recycling [14]. The ubiquitinylation of MET by the E3 ubiquitin ligase CBL (casitas B-lineage lymphoma), which depends on its recruitment to Y1003, marks the receptor for degradation through its inclusion in multivesicular bodies (MVB) and lysosomes [15]. Alternatively, MET can also be recycled via Rab11-positive recycling endosomes or vesicles containing GGA3 (Golgi-localized gamma ear-containing Arf-binding protein 3) [16].

Throughout the intracellular trafficking of MET, the phosphorylated receptor can activate different subsets of substrates depending on its endosomal location [14,17,18]. For instance, Kermorgant *et al.* demonstrated that STAT3 activation and nuclear accumulation are impaired upon HGF stimulation when MET internalization or its trafficking to perinuclear compartments is blocked [17]. On the other hand, although activation of the PI3K/AKT axis has been typically restricted to the plasma membrane, there is indication of HGF-induced AKT phosphorylation occurring post-endocytically [19]. These examples illustrate that a fine-tuned regulation of MET routing is essential in normal cells.

Among the proteins involved in the trafficking of internalized receptors are the members of the sorting nexins (SNX) protein family. These proteins are characterized by the presence of a PX (phox-homology) domain, allowing their association with endosomal membranes; other domains in SNXs are used for subclassification [20]. The presence of a BAR domain (Bin, Amphiphysin, Rvs) characterizes the heterodimer-forming SNX-BAR subfamily to which SNX1/2/5/6 belong [21]. SNX-BAR interaction and trafficking role on RTKs, such as EGFR, PDGFR, and IGF1R, have been reported [20–22]. Specifically, SNX2 is an interacting partner of MET [25,26]. Interestingly, low levels of *SNX2* mRNA and SNX1 protein in primary tumors have been correlated with a poorer prognosis in CRC patients [27,28]. Studies using siRNA-mediated down-regulation of SNX1 and SNX2 suggest their involvement in regulating MET signaling in lung cancer cells [26,29]. Furthermore, we have shown that SNX2 down-regulation enhanced MET-induced ERK1/2 (extracellular signal-regulated kinases 1 and 2) activation in HeLa cells [27]. However, the mechanisms underlying SNX1/2-controlled MET routing and signaling in CRC remain poorly understood. We hypothesized that SNX1 and SNX2 play a critical role in MET signaling, contributing to CRC progression. In this study, MET trafficking, signaling and regulated biological processes upon HGF stimulation were characterized following the knockout (KO) of *SNX1* and *SNX2* genes in the HCT116 CRC cell line.

Here, we observed reduced trafficking of MET to early endosomes in the *SNX1/2* KO cells, along with increased MET and AKT phosphorylation. Notably, the absence of both trafficking proteins boosted HGF-induced resistance to tumor necrosis factor-related apoptosis-inducing ligand (TRAIL)-driven apoptosis, without consequences on proliferation and migration.

## RESULTS

### Absence of SNX1 and SNX2 in HCT116 cells does not affect MET protein expression

To study the role of SNX1 and SNX2 in MET intracellular trafficking and signaling, we abrogated their expression in HCT116 CRC cells through CRISPR/Cas9 gene editing. SNX1/2 protein expression was validated by immunoblotting (**Figure 1A**) and genomic DNA sequencing of the CRISPR/Cas9 targeted regions showed biallelic frameshift nucleotide deletions (**Figure S1**). To explore potential compensation mechanisms, we evaluated protein expression of SNX5 and SNX6, both members of the SNX-BAR subfamily capable of forming heterodimers with SNX1/2 [21]. In *SNX1/2* KO cells, the presence of SNX5 and SNX6 was markedly decreased, but the level of VPS29, a key component of the retromer with which SNX1/2 interact, remained unchanged (**Figure 1A**). Given the established connection between the modulation of SNXs’ expression and the expression of membrane receptors [24,26,29,34], the level of MET at steady state was determined. *MET* mRNA was quantified by qRT-PCR in parental and *SNX1/2* KO cells (**Figure S2A**), revealing a decrease of close to 20% in the genetically modified cells. However, immunoblotting results demonstrated that MET protein level remained unchanged (**Figures 1B**), suggesting that SNX1/2 do not significantly impact MET protein expression in HCT116 cells.

**Figure 1:**
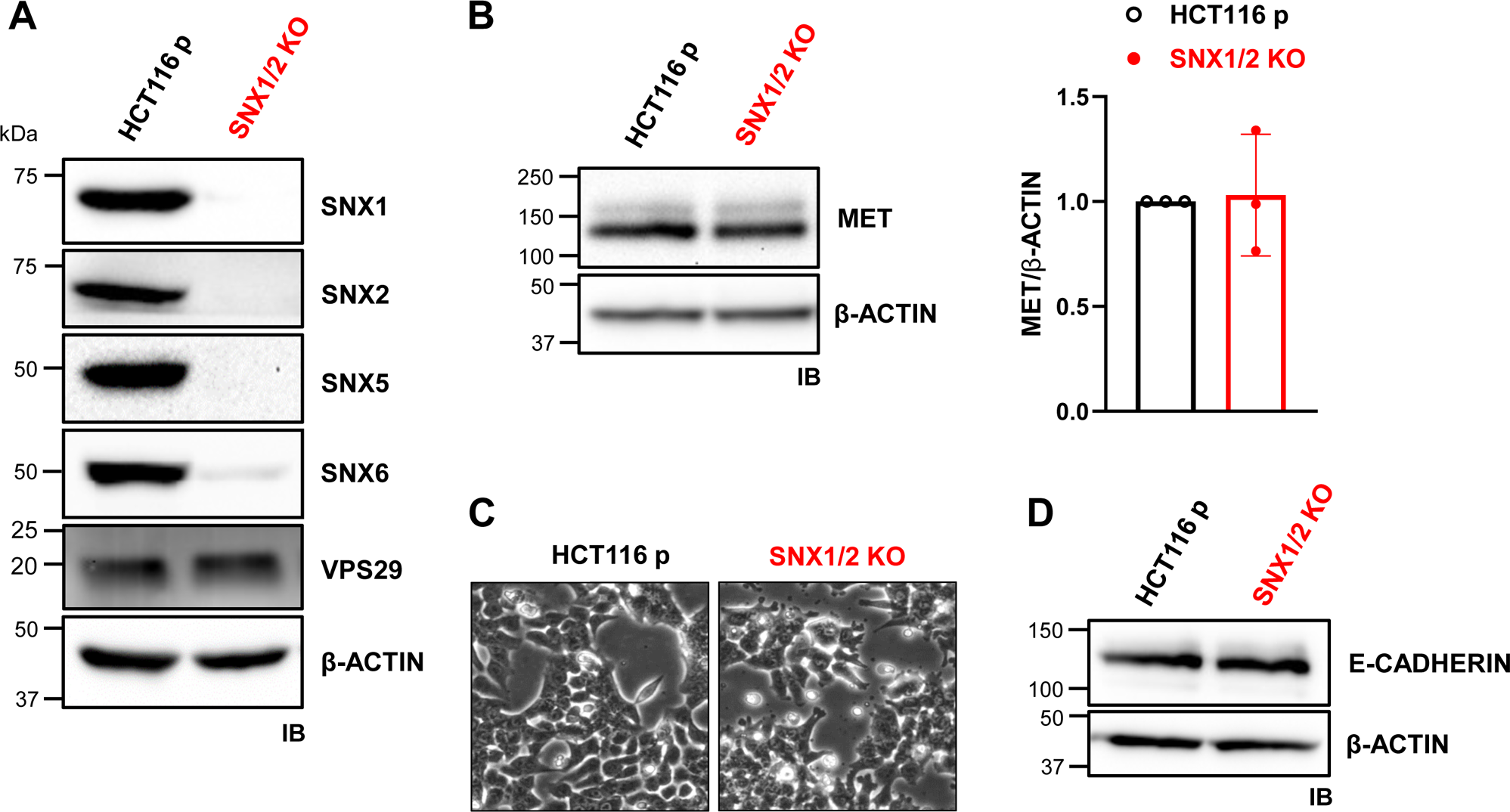
The knockout of *SNX1* and *SNX2* in HCT116 cells does not alter MET expression. **A, B** The KO of *SNX1* and *SNX2* was validated by immunoblotting as well as the expression level of SNX5, SNX6, VPS29 (**A**), and MET (**B**). Two bands corresponding to the precursor (170 kDa) and the mature receptor (145 kDa) MET receptor. Actin was used as a loading control. MET/β-actin ratio was quantified (N=3). **C** Morphology of parental and *SNX1/2* KO cells under phase-contrast microscopy. **D** E-cadherin protein expression under basal conditions was determined by immunoblotting.

We next determined if nullifying *SNX1* and *SNX2* altered the cells morphology. Phase-contrast microscopy analysis showed that parental and KO cells displayed a similar epithelial-like morphology, except for the presence of more abundant protrusions at the membrane of KO cells (**Figure 1C**). Staining of actin filaments with AlexaFluor488-labeled phalloidin revealed that these protrusions were rich in actin filaments, as well as the presence of a more diffuse cortical actin cytoskeleton in *SNX1/2* KO cells (**Figure S2B**). However, a similar level of E-cadherin was detected in parental and KO cells (**Figure 1D**). These observations suggest that the absence of SNX1 and SNX2 proteins has minimal impact on epithelial-to-mesenchymal transition (EMT) in HCT116 cells.

### Absence of SNX1 and SNX2 delays the entry of MET into early endosomes, but not its traffic to late endosomes

Following HGF binding, MET is rapidly internalized and localizes to early/sorting endosomes [18]. Given that SNX1/2 are located on early endosomes and are commonly involved in endosomal sorting of various receptors [24,26], we examined whether *SNX1/2* KO altered MET endocytic trafficking. First, we assessed the surface expression of MET in the absence of stimulation by flow cytometry. DsiRNA-mediated down-regulation of MET was used to confirm the specificity of the signal. Comparable levels of MET at the surface of parental (76.5% ± 0.7) and *SNX1/2* KO cells (75.3% ± 2.9) were observed (**Figure 2A**).

**Figure 2:**
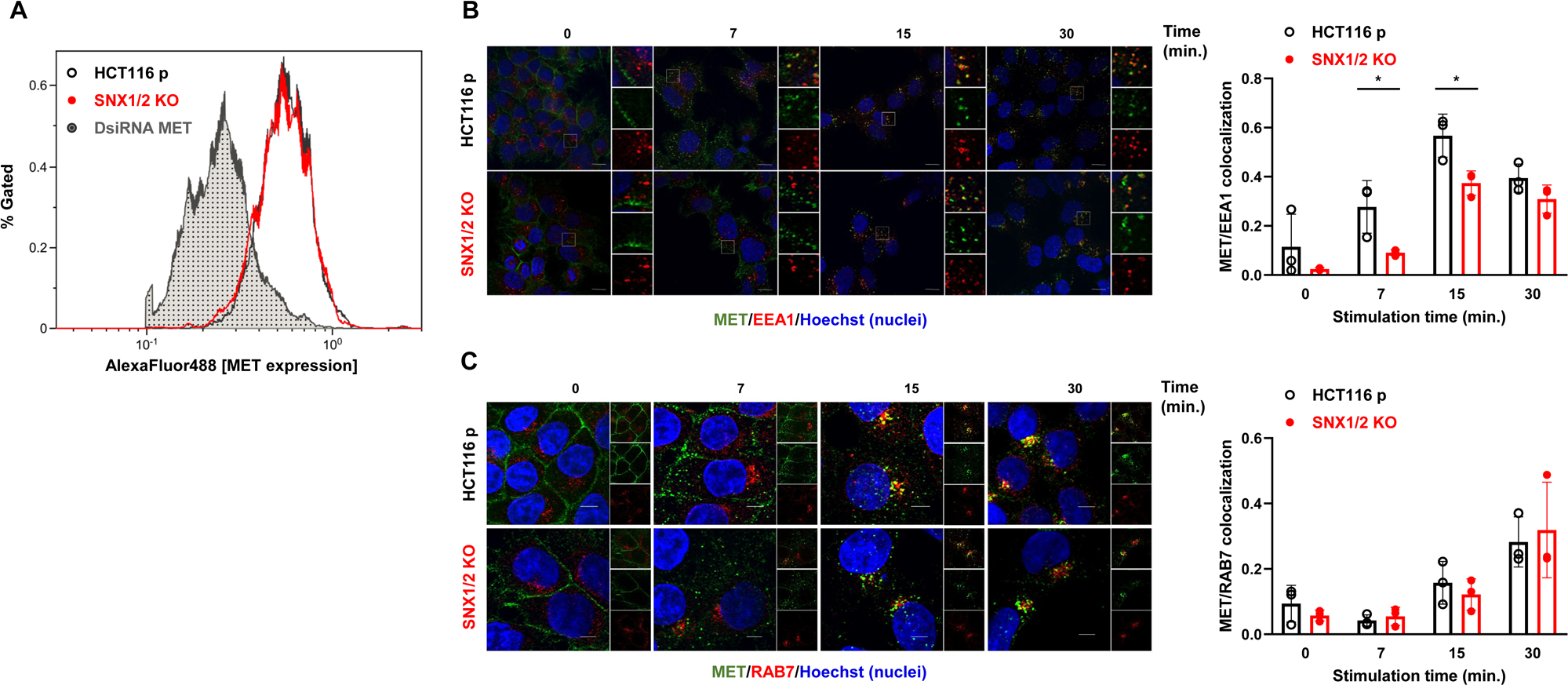
MET localization to early endosomes is decreased in *SNX1/2* KO HCT116 cells. **A** Cells were labeled with a MET-specific AlexaFluor488-conjugated antibody and the level of the receptor at the plasma membrane was determined by flow cytometry. Cells transfected with DsiRNA targeting MET were used as negative control. The histogram is representative of three independent experiments. **B, C** Confocal microscopy images of MET in parental and KO HCT116 cells stimulated with HGF (40 ng/ml) for the indicated period. Prior to stimulation, cell surface MET (green) was labeled with the anti-MET L6E7 antibody. Early or late endosomes were labeled (red) with EEA1 (**B**) or Rab7 (**C**), respectively. Nuclei were counterstained with Hoechst. Scale 10 μm; 60X objective. The bar graphs represent Mander’s colocalization coefficient ± SD (N=3).

Next, MET trafficking was examined following cell-surface receptor labeling and ligand-induced internalization. The presence of the receptor in the first step of the endocytic pathway was determined by co-labeling with the early endosomal marker EEA1 (early endosome antigen 1) [35], following 0, 7, 15 and 30 min of HGF stimulation (**Figure 2B**). The unstimulated receptor was located at the plasma membrane in parental and *SNX1/2* KO cells. Colocalization with early endosomes was observed after 7 min of HGF stimulation, reaching a maximum colocalization at 15 min in both parental and KO cells. Quantification of MET and EEA1 co-occurrence using Mander’s overlap coefficient revealed a significant reduction in MET localized with early endosomes at 7 and 15 min post-stimulation in *SNX1/2* KO cells compared to parental cells (*p* = 0.04 and *p* = 0.03, respectively).

We next examined whether MET trafficking to late endosomes was altered by assessing the level of MET colocalization with RAB7, a marker of late endosomes [36] (**Figure 2C**). The presence of MET in this compartment became apparent after 15 min and was higher after 30 min of stimulation. However, colocalization quantification did not show a significant difference between parental and *SNX1/2* KO cells. Together, these results indicate a possible role of SNX1 and SNX2 in early MET trafficking events.

### Absence of SNX1 and SNX2 does not affect MET degradation or recycling

To further determine the contribution of SNX1/2 on MET endosomal sorting, we studied the impact of the loss of SNX1/2 expression on MET degradation and recycling. The kinetic and extent of colocalization with LAMP1, a marker of the lysosomal compartment [37], was first used to analyze MET delivery to lysosomes. This analysis showed no significant difference between parental and *SNX1/2* KO cells (**Figure 3A**). This result was corroborated by immunoblotting of cells stimulated with HGF for a prolonged period in the presence of cycloheximide to prevent *de novo* protein synthesis (**Figure 3B**). The kinetic and level of MET degradation was not significantly different between parental and KO cells, indicating that ligand-induced lysosomal degradation of MET is not dependent on SNX1 and SNX2.

**Figure 3:**
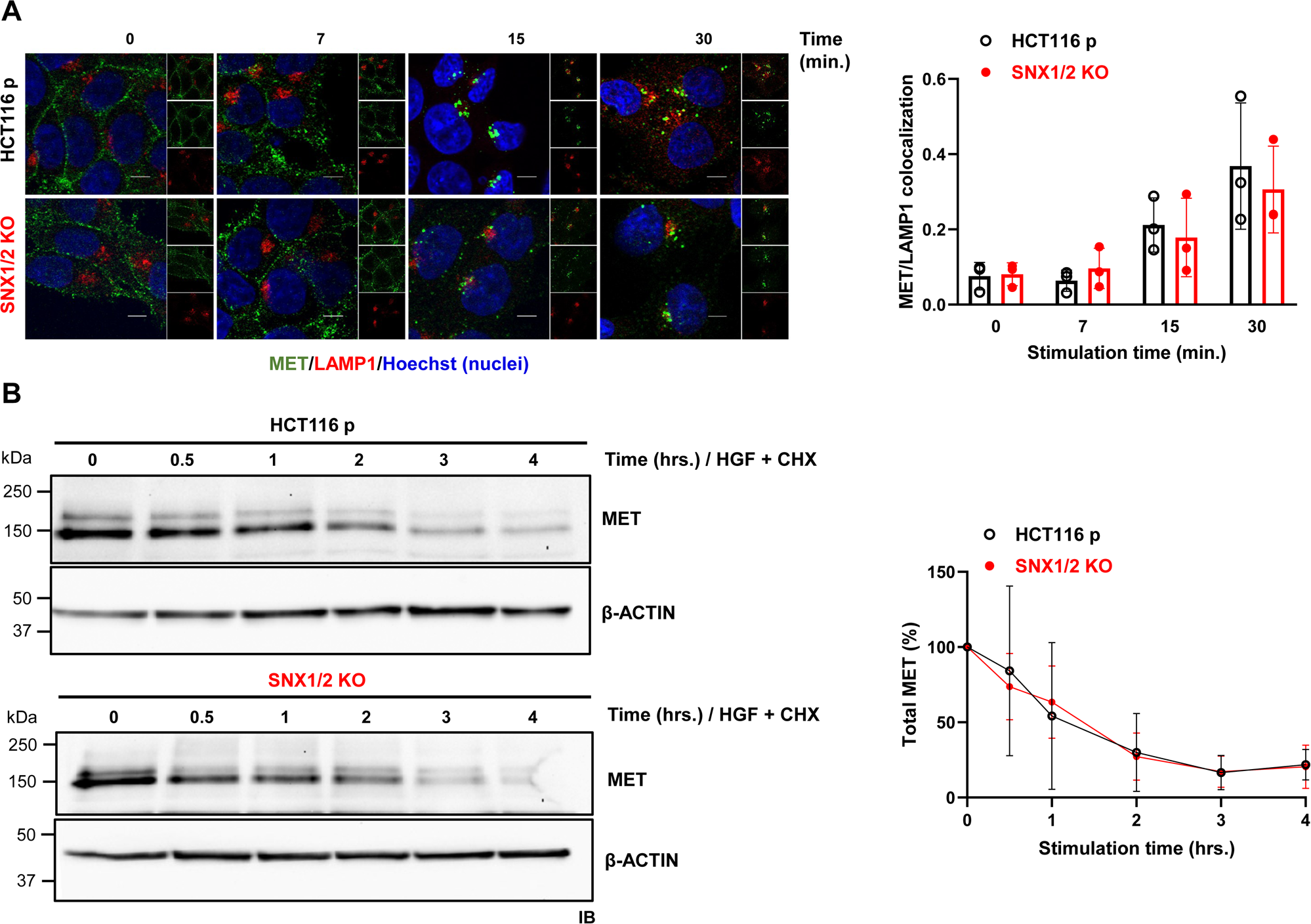
The knockout of *SNX1* and *SNX2* has no effect on MET degradation. **A** Confocal microscopy images of MET in parental and KO HCT116 cells stimulated with HGF (40ng/ml) for the indicated period. Prior to stimulation, cell surface MET (green) was labeled with the anti-MET L6E7 antibody. Lysosomes were labeled with Lamp1 (red). Nuclei were counterstained with Hoechst. Scale 10 μm; 60X objective. The bar graph represents Mander’s colocalization coefficient (N=3). **B** Expression level of MET was determined by immunoblotting from cell lysates obtained following HGF-stimulation of parental and *SNX1/2* KO HCT116 cells in the presence of cycloheximide (40 μg/mL) for the indicated time. Actin was used as a loading control. Graph shows the densitometric quantification of MET protein normalized to the unstimulated sample (N=3).

Besides being directed to the lysosomes, a fraction of activated MET undergoes either rapid or slow recycling back to the plasma membrane [38]. Both slow and fast MET recycling were evaluated by colocalization with CD71 positive recycling endosomes [39] or GGA3-containing vesicles [16], respectively. MET was mainly located in both compartments following 15 min of HGF stimulation (**Figure 4**). Quantification of Mander’s coefficient did not reveal significant differences between parental and *SNX1/2* KO cells. These findings indicate that SNX1 and SNX2 are not required for MET recycling and suggest that the delay of MET localization in early endosomes induced by the loss of SNX1/2 is not attributed to an increased MET recycling.

**Figure 4:**
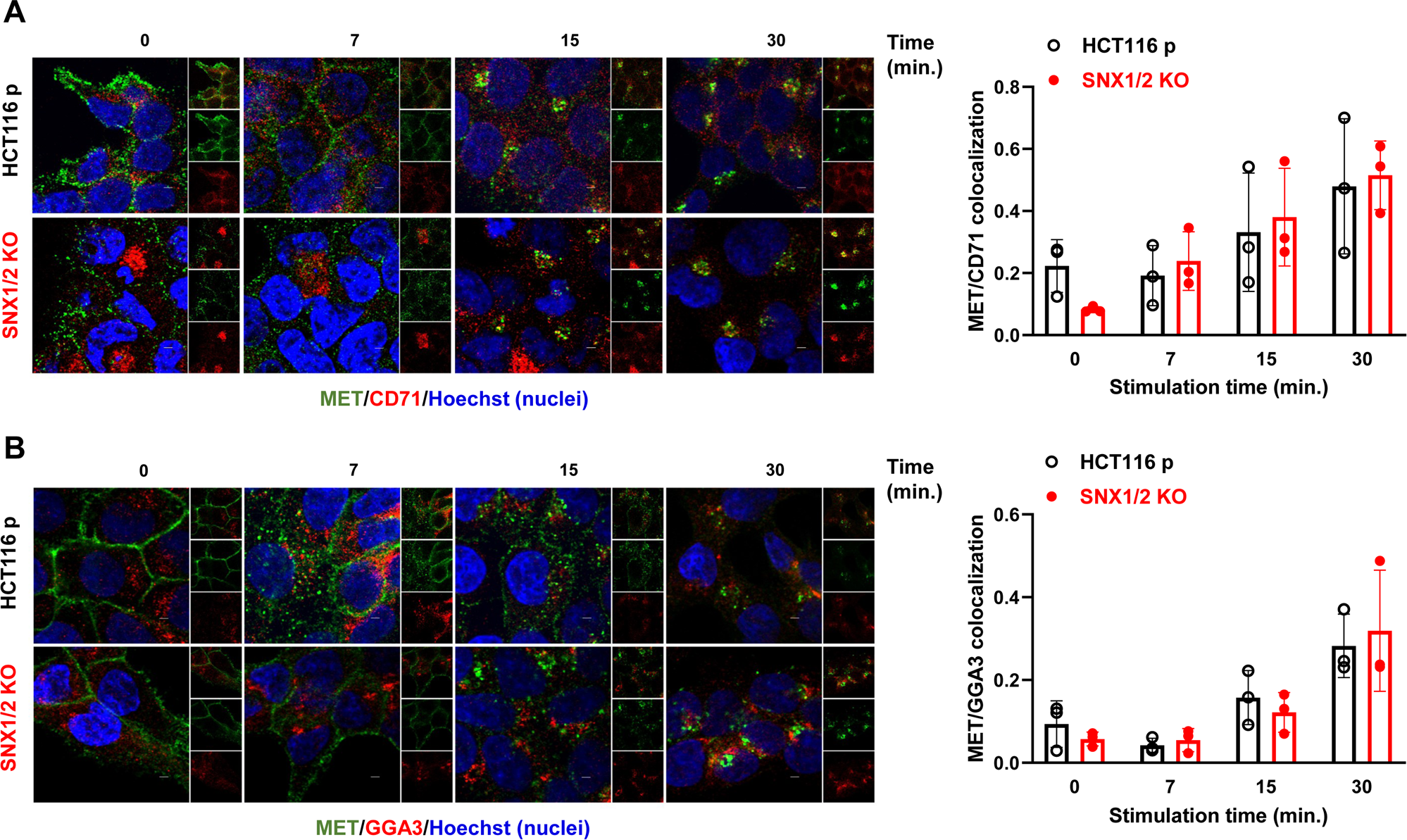
MET recycling is not altered in *SNX1/2* KO HCT116 cells. Confocal microscopy images of MET in parental and KO HCT116 cells stimulated with HGF (40 ng/ml) for the indicated period. Prior to stimulation, cell surface MET (green) were labeled with the anti-MET L6E7 antibody. Slow and fast recycling endosomes were respectively labeled (red) with CD71 (**A**) or GGA3 (**B**). Nuclei were counterstained with Hoechst. Scale 10 μm; 60X objective. The bar graphs represent Mander’s colocalization coefficient (N=3).

### Absence of SNX1 and SNX2 enhances HGF-induced MET and AKT phosphorylation

The consequence of the loss of SNX1/2 protein expression on HGF-mediated signaling was next investigated. For this, the phosphorylation status of MET and downstream effectors following ligand stimulation of serum-starved HCT116 cells was evaluated through immunoblotting. Stimulation with HGF resulted in rapid MET phosphorylation, detectable as early as 7 min, in both parental and *SNX1/2* KO cells (**Figure 5**). However, in the KO cells, this response was enhanced, displaying a delay in reaching the activation peak, whereas MET phosphorylation levels were decreased to a similar extent in both cells at 60 min. Importantly, the increase in MET phosphorylation in *SNX1/2* KO cells occurred despite similar levels of the total receptor (**Figure S3A**).

**Figure 5:**
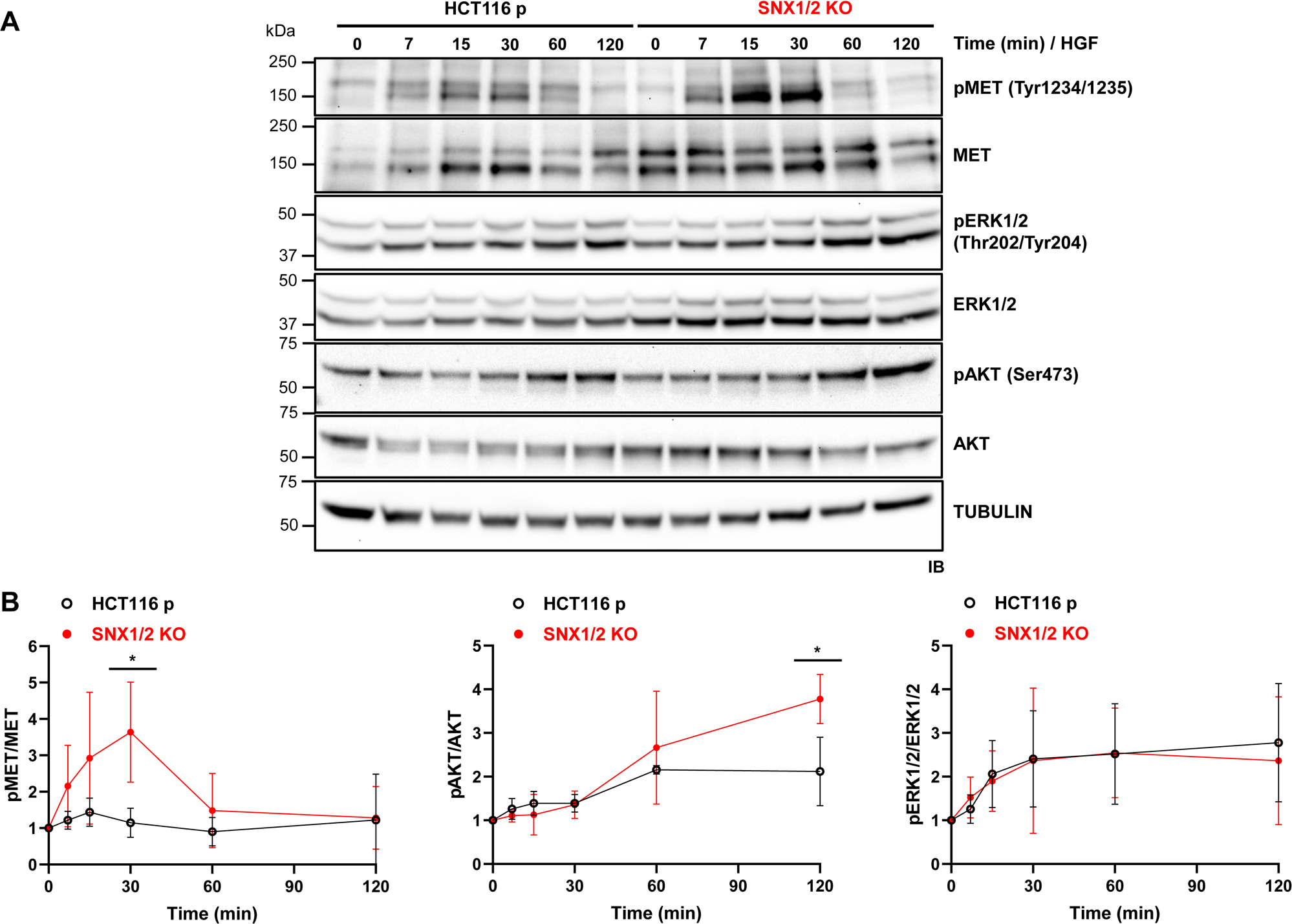
The knockout of *SNX1* and *SNX2* potentiates HGF-induced MET and AKT phosphorylation. **A** Serum-starved cells were stimulated with 50 ng/mL HGF for the indicated period. Phosphorylation of MET and downstream effectors AKT and ERK1/2 was determined by immunoblotting. Tubulin was used as a loading control. **B** Graphs show the densitometric quantification of data from **A** expressed as the ratio of phosphorylated protein/total protein and normalized to the unstimulated sample. Data correspond to the mean ± SD (N=3).

Stimulation of MET by HGF triggers, among others, the activation of the Ras/MAPK and PI3K/AKT pathways [12]. In HGF-stimulated cells, ERK1/2 phosphorylation showed a time-dependent increase, remaining constant after 30 min, but it was not significantly different between parental and *SNX1/2* KO cells (**Figure 5**). However, AKT showed increased phosphorylation at later time points (60 and 120 min) in KO compared to parental cells. Importantly, total AKT and ERK1/2 protein levels remained unaffected upon HGF stimulation in both cell lines (**Figures S3B, S3C**). Notably, the kinetics of HGF-induced MET and AKT phosphorylation observed in *SNX1/2* KO cells was not an artifact of clonal variation, as similar results were obtained in HCT116 cells in which SNX1/2 expression was transiently down-regulated by DsiRNA transfection (**Figure S4**).

The observed HGF-induced ERK1/2 and AKT phosphorylation in the absence or down-regulation of SNX1/2 was relatively modest, despite the robust activation of MET. Considering that HCT116 cells harbor *KRAS* and *PI3K* mutations, resulting in their constitutive activation [40], we considered the possibility of a negative feedback mechanism acting in Ras/MAPK and PI3K/AKT pathways in these cells. To address this hypothesis, serum-starved cells were treated with a strong stimulus (FBS) and the activation of ERK1/2 and AKT was analyzed. Under these conditions, the phosphorylation status of ERK1/2 slightly increased, and that of AKT remained unchanged in parental cells upon FBS stimulation (**Figure 6**). However, in the KO cells a similar pattern was exhibited by ERK1/2, while AKT phosphorylation was significantly boosted. Altogether, these results suggest that the loss of SNX1/2 might disrupt a negative regulatory mechanism in the PI3K/AKT axis.

**Figure 6:**
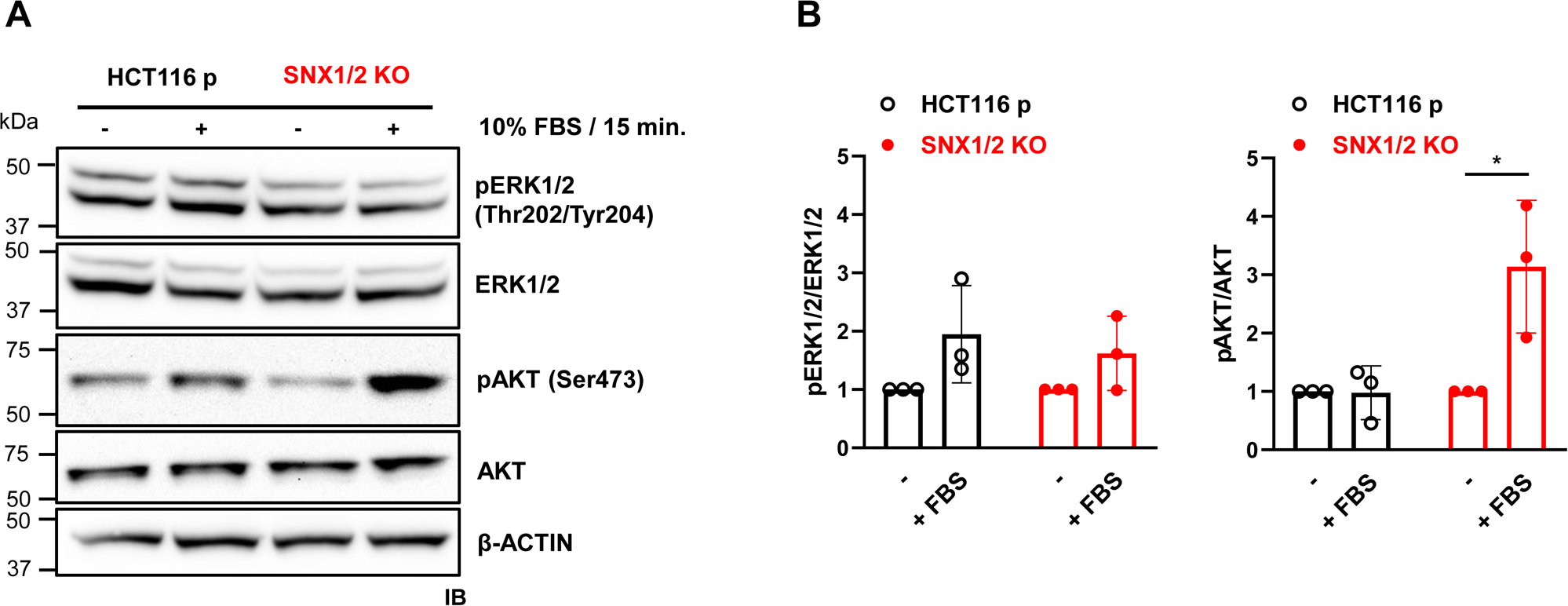
The knockout of *SNX1* and *SNX2* reverses negative feedback regulation of the PI3K-AKT axis. **A** Serum-starved cells were stimulated with 10% FBS for 15 min. Phosphorylation of ERK1/2 and AKT was determined by immunoblotting. Actin was used as a loading control. **B** The bar graphs show the densitometric quantification of data from **A** expressed as the ratio of phosphorylated protein/total protein normalized to the unstimulated cells. Data correspond to the mean ± SD (N=3).

### Absence of SNX1 and SNX2 does not affect HGF-induced proliferation or migration

Cellular responses ascribed to HGF stimulation include morphogenesis, which implicate the coordination among others, of proliferation, migration, and invasion [41]. Given the potentiated MET signaling in *SNX1/2* KO cells, we explored some of these responses. We first assessed the impact of *SNX1/2* loss on cell growth under normal culture conditions in the presence of FBS. We found that both parental and KO cells grew at similar rates, with doubling times of 25.5 and 24 h, respectively (**Figure 7A**). The specific contribution of HGF to cellular proliferation was also determined by conducting EdU incorporation assays under serum starvation and upon MET stimulation. The latter caused a 2-fold increase in the number of proliferative cells, with no apparent difference observed for *SNX1/2* KO cells (**Figure 7B**).

**Figure 7:**
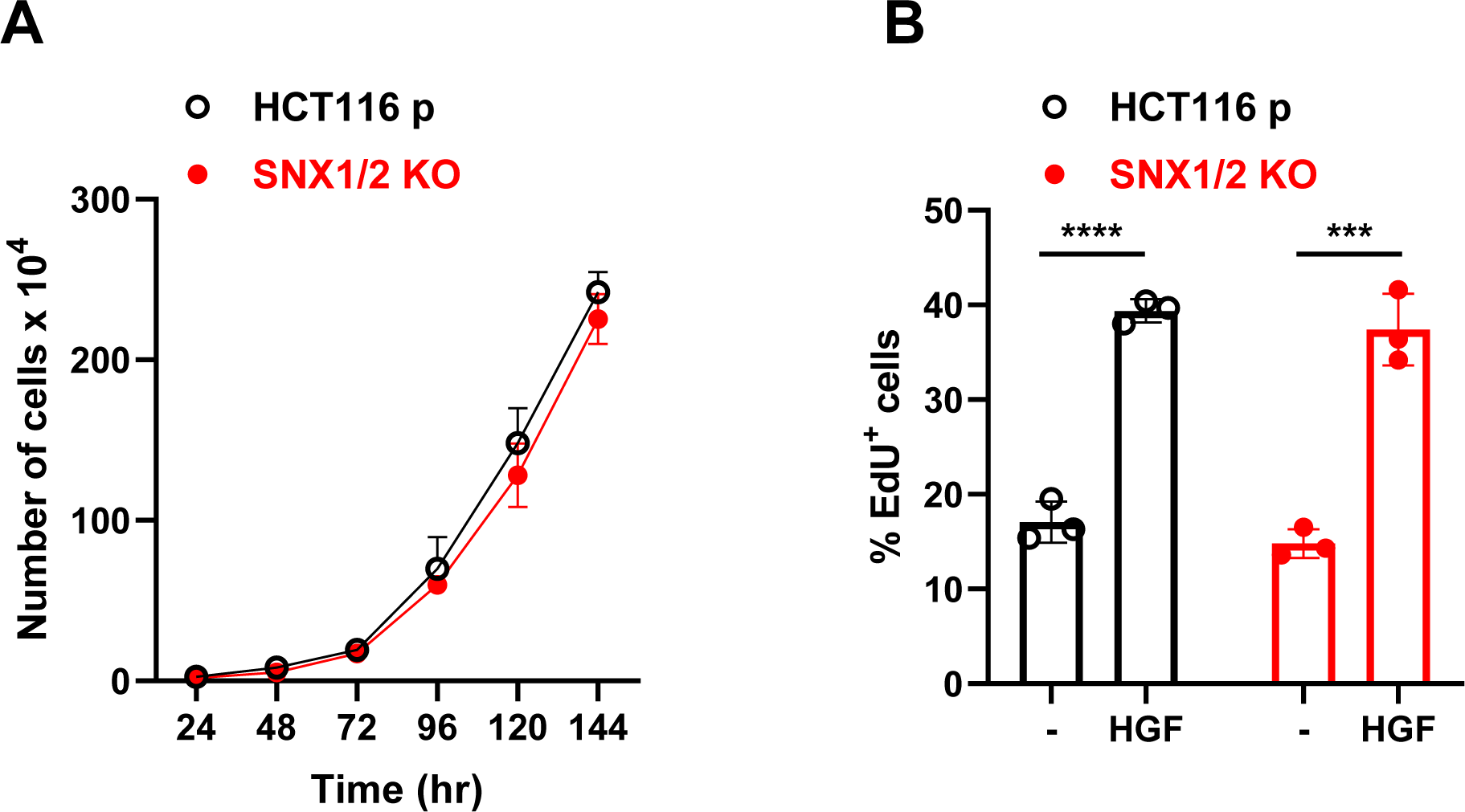
The proliferation of HCT116 cells was not affected by *SNX1* and *SNX2* KO. **A** Representative growth curves of parental and *SNX1/2* KO HCT116 cells. Calculated doubling time was calculated as 25.5 ± 3.6 h (95% CI) and 24 ± 2.9 h for parental and KO cells, respectively (N=3). **B** The graph shows the percentage of 5-ethynyl-2’-deoxyuridine (EdU) positive cells following stimulation with HGF (50 ng/mL) for 12 hours. Data correspond to the mean ± SD (N=3).

We also investigated if SNX1/2 loss influenced motogenic responses by performing cell scattering and wound healing assays. Phase-contrast images showed a clustered cell morphology in serum-deprived parental, and particularly noticeable in *SNX1/2* KO cells (**Figure 8A**). Nevertheless, following HGF stimulation, both cells scattered similarly. Moreover, both the basal and HGF-induced migration capacities of parental and *SNX1/2* KO cells were similar, as assessed in a wound healing assay. (**Figure 8B and C**). Even though HGF allowed faster gap closure after 24 h, no differences in the rate of wound closure were detected between parental and *SNX1/2* KO cells (**Figure 8C**). Therefore, the loss of SNX1/2, despite leading to increased HGF-induced MET signaling, does not influence cell proliferation or migration.

**Figure 8:**
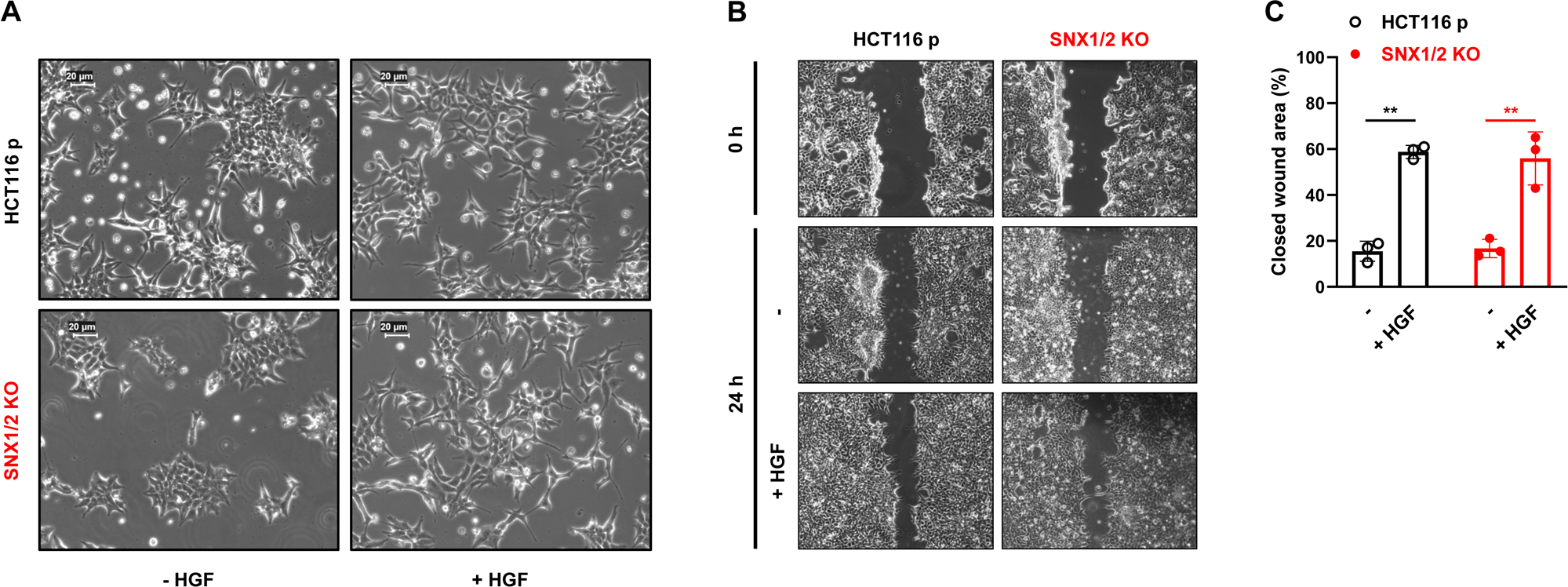
*SNX1* and *SNX2* KO in HCT116 cells has no effect on HGF-stimulated induced scattering and migration. **A** Scatter activity of HGF (50 ng/mL; 24 hours) on parental and *SNX1/2* KO cells. **B** Confluent cell monolayers were scratched, and the ability to fill the gap was monitored in the presence or not of HGF (100 ng/mL) after 24 h. **C** Bar graph shows the mean percentage of wound healing ± SEM (N=3).

### Absence of SNX1 and SNX2 increases HGF-driven protective effect from TRAIL-induced apoptosis

Activation of the HGF/MET receptor axis protects against multiple cell death-inducing challenges [12]. Thus, we assessed if the absence of SNX1/2 influenced cell response to TRAIL-induced apoptosis in the presence or not of HGF stimulation. Cells were concomitantly treated with TRAIL (50 ng/mL) and HGF (100 ng/mL) for 2 h, and the extent of apoptosis was estimated by PARP-1 cleavage, a robust hallmark of this type of cell death [42].

Under TRAIL treatment, a band corresponding to the cleaved form of PARP-1 (89 kDa) was detected, concomitant with the disappearance of its full-length form, in both parental and *SNX1/2* KO cells (**Figure 9A**). The ratio of cleaved to uncleaved (full-length) PARP-1 was quantified as a measure of the apoptotic response. This ratio indicated that the sensitivity of parental and *SNX1/2* KO cells to TRAIL-induced apoptosis was similar in the absence of HGF (**Figure 9B**). However, the protective effect of HGF stimulation against TRAIL-induced apoptosis was significantly increased in the *SNX1/2* KO cells (**Figure 9B**). Similarly, HGF-induced AKT phosphorylation was enhanced in TRAIL-treated *SNX1/2* KO cells (**Figure 9C**), and treatment with the PI3K inhibitor LY294002 blunted both AKT phosphorylation and the anti-apoptotic effect conferred by HGF stimulation, as expected (**Figure S5**). These results indicate that the HGF-mediated anti-apoptotic response against TRAIL-induced apoptosis takes place via the PI3K/AKT axis-dependent pathway in HCT116 cells.

**Figure 9:**
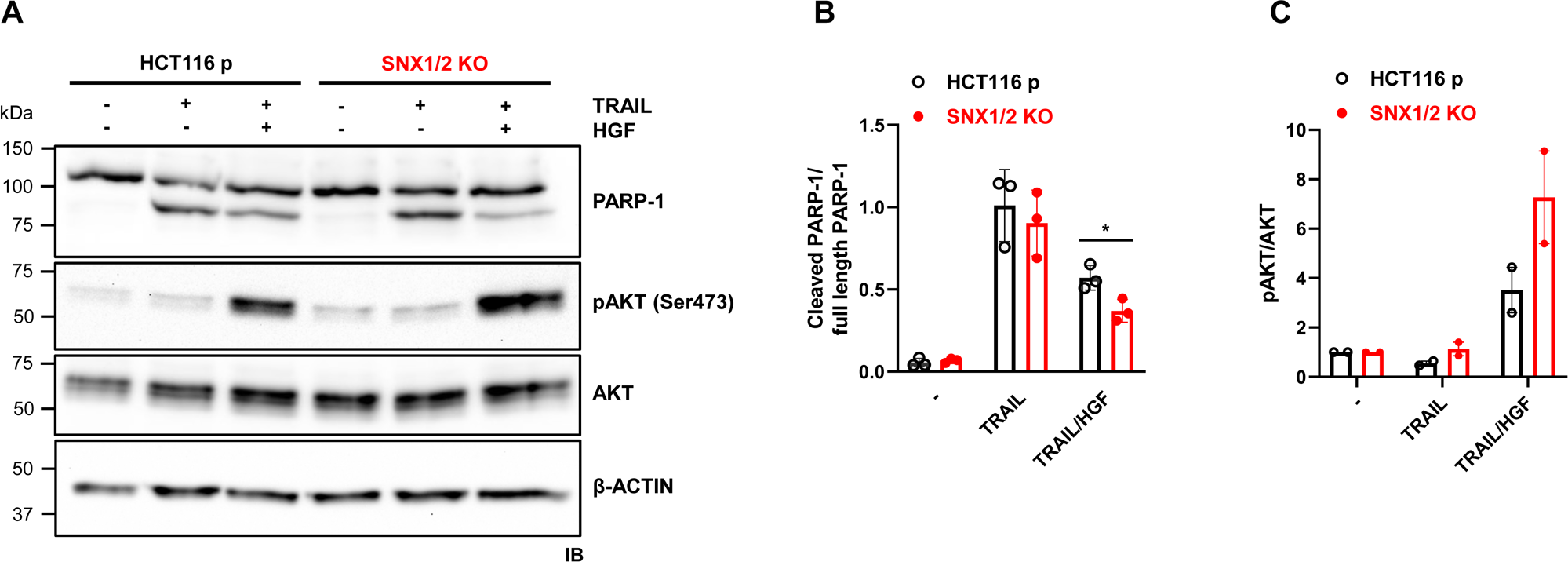
HGF-dependent protective effect from TRAIL-induced apoptosis in *SNX1/2* KO HCT116 cells. **A** Apoptosis was induced by treating cells with TRAIL (50 ng/mL) alone or in combination with HGF (100 ng/mL). The cleavage of PARP-1, along with the total and phosphorylation levels of AKT protein, was determined by immunoblotting, with actin used as a loading control. Bar graphs show the densitometric quantification of **B** full-length PARP-1 (left) or cleaved/intact ratio (right) and **C** the ratio of phosphorylated AKT/total AKT, and normalized to the unstimulated sample, for each condition. Data correspond to the mean ± SD (N=3).

## DISCUSSION

As part of the endosomal trafficking machinery, the family of SNX proteins is involved in the sorting of several types of receptors [26,34,43,44]. Particularly, SNX1/2 have been reported as regulators of EGFR routing to the degradative pathway [24,34,43]. Despite a limited knowledge of a link between SNX and MET, a constitutive and direct interaction between SNX2 and MET has been identified through two-hybrid screening [25], and a study demonstrated that silencing SNX2 in lung cancer EBC-1 cells led to reduced MET protein levels and phosphorylation, contributing to overcoming resistance to anti-EGFR drugs [26]. As only fragmentary evidence exists, the role of SNX1/2 in MET trafficking and signaling, and how changes in their expression might contribute to malignancy in CRC is poorly understood. In this study, we showed that SNX1/2 participate in the efficient trafficking of MET toward early endosomes, thereby influencing the level of activation of the receptor and AKT, as well as cell sensitivity to apoptosis.

Our results indicate that the complete abrogation of SNX1/2 expression in HCT116 cells neither interferes with the total levels nor with the membrane expression of MET at a steady state. This agrees with our previous work showing no alteration in MET levels when SNX2 expression was reduced in HeLa cells [27]. However, studies have reported that down-regulation of SNX1 or SNX2 is associated with a decrease in total MET levels in lung cancer cells, while another study has reported that a decrease in SNX1 reduces cell membrane presence and leads to a concomitant increase in intracellular compartments’ distribution of the receptor in A549 lung cancer cells [26,29]. Discrepancies with our results could be explained by the fact that others have depleted only one SNX and/or used different cells lines. By down-regulating SNX1/2, we found a simultaneous decrease in SNX5/6 expression in HCT116 cells. These four proteins act together in different dimer combinations [21], mainly in association with the retromer [45] or as part of the ESCPE-1 complex (endosomal SNX-BAR sorting complex for promoting exit) [44]. We surmise that the decrease in SNX5/6 expression is attributed to the absence of their SNX1/2 dimerization partners, a phenomenon often seen for members of multiprotein complexes [46]. Thus, any effect we observed is a combination of SNX1/2 absence and SNX5/6 reduction.

In our studies, the lack or decrease in SNX1/2/5/6 did not cause a drastic change in the morphology of HCT116 cells, as analyzed as part of the phenotypical cellular characterization. Certain changes in cellular morphology ascribed to the EMT process are frequently considered clues of a malignant phenotype, such as the acquisition of fibroblastic-like morphology, cytoskeleton rearrangement, and breakdown of cell-cell junctions [47]. We only observed more abundant membrane protrusions in the KO cells, which could be the result of actin reorganization. Others have demonstrated that SNX1 overexpression contributes to a reversion of EMT in gastric cancer cells [48], while the knockdown of SNX5 in clear-cell renal-cell carcinoma cells promotes it [49], supported by changes in the expression of EMT markers such as Vimentin, Snail, E-cadherin, Claudin-1, or ZO-1 (zonula occludens-1). In our study, although other EMT markers remain to be evaluated, unchanged E-cadherin expression levels after depleting SNX1/2 indicates that the observed morphological changes might not precisely be accompanied by EMT.

Upon HGF stimulation, MET is internalized in clathrin-coated vesicles and then delivered to early endosomes from which the receptor is sorted to late endosomes/lysosomes for degradation or recycled back to the plasma membrane [16,50]. In *SNX1/2* KO cells, a significant decrease in colocalization between MET and the early endosomal marker EEA1 was observed at early time points, but not at 30 min post HGF stimulation. Furthermore, no significant difference was observed in MET trafficking to late endosomes (RAB7), lysosomes (LAMP1), or recycling endosomes (CD71, GGA3). The observed delay in the entry of MET into early endosomes in the KO cells suggests that SNX1/2 could mediate early trafficking steps of MET. However, the fact that this did not result in a delay of MET trafficking throughout the following endocytic compartments is intriguing. We hypothesize that MET undergoes endocytosis through two parallel pathways. One pathway delivers the receptor directly to EEA1-labeled endosomes, while the other passes through an intermediate (EEA1-negative endosome) before reaching the EEA1-labeled endosome [51]. In this paradigm, the absence of SNX1/2 redirects receptor trafficking to this second endocytic pathway, explaining the delayed appearance of MET in EEA1-labeled endosomes without impacting the subsequent kinetics within the endocytic pathway. The nature of this second endocytic pathway remains unclear but studies on GPCRs [52] and EGFR [53,54] point to their internalization in a very early endosomal (VEE) sub-compartment devoid of EEA1 markers but positive for APPL1 (adaptor protein, phosphotyrosine interacting with pleckstrin homology (PH) domain and leucine zipper 1). APPL1 is a multiprotein adaptor containing a PH and a PTB (phospho-tyrosine binding) domain, in addition to a structured N-terminal BAR domain [55], like SNX-BAR proteins. Of note, the PH and PTB domains are found in many proteins that are part of the MET signalosome, like AKT, GAB1/2, and SHC proteins [56,57]. The APPL1-labeled endosomes have been demonstrated to mature into EEA1-positive endosomes [58,59] and play a specialized role in trafficking cargo devoted to signal transduction. Specifically, APPL1 interacts with several receptors and components of signaling pathways, including AKT [60–62]. While an interaction between APPL1 and MET has not been described yet, studies suggest that HGF-dependent AKT activation requires APPL1 for deploying survival and migration responses in murine fibroblasts [19]. The subcellular location of APPL1 proteins implies that this AKT signaling takes place in VEEs. Therefore, we propose that suppressing SNX1/2 expression redirects the routing of MET into VEE and/or retards VEE maturation into EEA1-endosomes, prolonging the retention time of MET in VEE and enhancing MET signaling. This hypothesis will be validated in future studies.

Our present results showed a boosted MET and AKT phosphorylation, but not that of ERK1/2 in *SNX1/2* KO HCT116 cells. Previously, we had reported an increase in MET and ERK1/2 phosphorylation following HGF stimulation in SNX2 knockdown HeLa cells [27]. We explain this discrepancy by differences between HeLa and HCT116 cells in the mutation status of upstream activators of AKT and ERK1/2. We propose that the phosphorylation pattern followed by AKT and ERK1/2 was influenced in HCT116 cells by the presence of activating mutations in *PI3K* (H1047R) and *KRAS* (G13D) [40], that are absent in HeLa cells [63]. The existence of such mutations in HCT116 likely could have contributed to the excessive activation of ERK1/2 and AKT, triggering negative feedback mechanisms. These regulatory controls in tumoral cells prevent over-activation of certain signaling pathways that could otherwise lead to senescence [64]. The modest increase in AKT and ERK1/2 phosphorylation following an acute FBS stimulus (**Figure 6**) indicates that negative feedback mechanisms are acting in PI3K/AKT and Ras/ERK1/2 pathways in parental HCT116 cells. Intriguingly, in the absence of *SNX1* and *SNX2*, AKT phosphorylation heightens upon HGF and FBS stimulation, suggesting a relief of negative feedback [65] in the PI3K/AKT pathway in KO cells. It remains to be elucidated which regulatory mechanism of the PI3K/AKT pathway is affected by the loss of SNX1/2, but we speculate that the altered MET and probably AKT localization could have interfered with the coordination of the molecular actors of feedback loops.

The activation of MET can evoke, among others, proliferation, scattering, and migration [12]. We observed that HGF potentiated all these behaviors, but that SNX1/2 do not seem to be determining elements. The development of such complex responses implicates the convergence of several signaling pathways, with differential contribution. For example, initial studies on MET receptor indicated that its stimulation promotes cell proliferation mainly via Ras/MAPK, with a role also attributed to the PI3K axis [12]. Scattering and migration require the implication of PI3K [66], which simultaneously with Ras/MAPK and STAT3 activation facilitates the induction of the tumoral invasive program [13]. Through PI3K, HGF-induced scattering and migration are mediated by the activation of the small GTPase Rac1, but they are AKT-independent [67]. This aligns with the observed similarity in these motility responses between parental and KO cells. In *SNX1/2* KO cells, only MET and subsequent AKT activation were enhanced after HGF stimulation. This could explain why we did not obtain differences in responses that are mainly associated with Ras/MAPK, such as proliferation.

We also verified resistance to apoptosis, which is frequently mediated by the PI3K/AKT pathway. We showed that the absence of SNX1/2 diminished the proteolysis of the hallmark protein PARP-1 during TRAIL-induced apoptosis, exclusively upon MET stimulation, which reveals increased resistance to cell death. PARP-1 is a caspase-3/7 substrate [68,69] and its cleavage depends on the full activation of the apoptotic cascade, which is regulated at several upstream steps including by RTK signaling. For instance, activated AKT can phosphorylate caspase-9, blocking the activation of this initiator caspase and downstream executioner caspase-3/7 [70]; engagement of the intrinsic caspase-9-driven apoptotic pathway is required for TRAIL-induced apoptosis in HCT116 cells [72]. Our results suggest that the potentiated MET-AKT activation exhibited by *SNX1/2* KO cells could be determining the reduction in PARP-1 cleavage in these cells, as a readout of their survival capacity.

In conclusion, our study demonstrates that SNX1 and SNX2 participate in initial endocytic trafficking steps that deliver MET to early endosomes in HCT116 CRC cells, consequently regulating MET signaling via PI3K/AKT and ultimately impacting cell survival. Besides confirming the intricate relationship between receptor trafficking and signaling, our findings might contribute to explain SNX1 and SNX2 down-regulation in CRC as an element disturbing MET activation in favor of malignancy [27,28]. The relevance of MET receptor in CRC progression [10] justifies the need of deeply understanding the mechanisms that lead to signaling deregulation. New regards on cell signaling go beyond the established model of MET endocytosis as a means of signal abrogation. Indeed, signals emanating from diverse intracellular locations have been uncovered and linked to specific biological effects [14,17,18]. This study opens new avenues for the development of innovative treatments for CRC, emphasizing the potential therapeutic implications of targeting MET sorting, particularly through interventions involving SNX proteins [73].

## MATERIALS AND METHODS

### Antibodies and reagents

Antibodies are listed in **Supplementary Table S1**. Recombinant human soluble TRAIL (ALX-201-073) was purchased from Enzo Life Sciences (Farmingdale, NY, USA). Human HGF (100-39) was from PeproTech (Cranbury, NJ, USA), and the PI3K inhibitor LY294002 was obtained from Cell Signaling Technology (Danvers, MA, USA). Cycloheximide (CHX; 239764) was from Sigma-Aldrich (St-Louis, MO, USA). General chemicals were from Sigma-Aldrich or Thermo Fisher Scientific (Waltham, MA, USA).

### Cell culture, transfection, and treatments

Human HCT116 colorectal carcinoma cell line (CCL-247) was purchased from ATCC (Cedarlane, Burlington, ON, Canada). Cells were maintained in Dulbecco’s modification Eagle’s medium (DMEM) supplemented with 10% fetal bovine serum (FBS), 2 mM L-glutamine, and penicillin-streptomycin antibiotics (Wisent Inc., Saint-Bruno, QC, Canada) in a 5% CO_2_ atmosphere at 37°C. The *SNX1/2* KO cells were generated by CRISPR/Cas9 gene-editing. The U6-gRNA/CMV-eCas9-2a-tGFP vector (Sigma-Aldrich) containing the guide RNA targeting *SNX1* (GGACAACACGGCATTGTCA) or *SNX2* (AAACGATTCGAAAAGAAGT) was transfected using Lipofectamine 2000 following the manufacturer’s instructions (Thermo Fisher Scientific). After 48 h, GFP-positive cells were selected and seeded individually in 96-well plates by flow cytometry (BD FACSAria^TM^ III, BD Biosciences, Franklin Lakes, NJ, USA). Double *SNX1* and *SNX2* KO cells were obtained using *SNX2* KO cells. *SNX1* and *SNX2* abrogation was verified by immunoblotting and genomic DNA sequencing. For experiments, cells were serum-starved for 16 h and exposed to one or to a combination of the following reagents: TRAIL (50 ng/mL), CHX (40 μg/mL), LY294002 (10 µM) and HGF (50 ng/mL or 100 ng/mL).

### Immunoblotting

Protein cell lysates were obtained by washing cells with ice-cold PBS (phosphate buffered saline; 123 mM NaCl, 10.4 mM Na_2_HPO_4_, 3 mM KH_2_PO_4_, pH 7.4) and collecting them using a cell scraper. For apoptotic assays, detached cells were also collected. Cell pellets were obtained by centrifugation at 2,000 x g for 5 min and lysed for 30 min on ice in modified RIPA buffer (radio-immunoprecipitation assay, 50 mM Tris, pH 7.4, 100 mM NaCl, 1% Nonidet P-40, 0.5% sodium deoxycholate, and 0.1% SDS) or Triton X-100 buffer (50 mM HEPES, pH 7.4, 150 mM NaCl, 0.5% Triton X-100, 10% glycerol, 2 mM EGTA, and 1.5 mM MgCl_2_) for MET immunoblotting. In both cases, protease and phosphatase inhibitors were added (1 mM 1,10-orthophenanthroline, 1 mM Na_3_VO_4_, 1 mM NaF, 50 µM 3,4-dichloroisocoumarine, 10 µM E64, and 10 µM leupeptin). Lysates were clarified at 18,000 x g for 15 min and protein concentration was determined by the bicinchoninic acid method (BCA; Thermo Fisher Scientific). Protein samples were separated on ammediol SDS-PAGE [30] and transferred to a PVDF membrane (polyvinylidene difluoride, EMD Millipore, Burlington, MA, USA) in transfer buffer (10 mM N-cyclohexyl-3-aminopropanesulfonic acid, pH 11, 10% methanol). Membranes were blocked with PBS containing 0.1% (v/v) Tween-20 and either 5% (w/v) non-fat dry milk (Carnation) or 3% bovine serum albumin (BSA, Sigma-Aldrich) for phosphorylated proteins detection. Ensuing membranes were incubated overnight at 4°C with primary antibodies followed by 1 h incubation with HRP (horseradish peroxidase)-conjugated secondary antibodies. Immune complexes were visualized by chemiluminescence using Immobilon Crescendo Western HRP substrate (EMD Millipore) or Clarity Max Western ECL Substrate (Bio-Rad, Hercules, CA, USA). Images were acquired using a VersaDoc 4000mp imager (Bio-Rad) and densitometric analyses were done using the QuantityOne software v4.6.7 (Bio-Rad). The membranes were stripped with stripping buffer (62.5 mM Tris, pH 6.8, 2% SDS, and 100 mM β-mercaptoethanol) under agitation at 50°C for 30 min and subsequently re-probed with a different primary antibody.

### Immunofluorescence

Cells were seeded on poly-L-lysine-coated (Sigma-Aldrich) glass coverslips (ThermoFisher Scientific). After 48 h, cells were serum-starved for 1 h in DMEM supplemented with 0.2 % (w/v) BSA and 25 mM HEPES. Cell surface MET were labeled with the anti-MET L6E7 antibody for 1 h at 4°C, followed by HGF stimulation for different periods. After stimulation, cells were fixed for 30 min with 3% (w/v) paraformaldehyde (Electron Microscopy Sciences, Hatfield, PA, USA), permeabilized for 10 min with 0.1 % Triton X-100 and blocked for 30 min with 10% goat serum before incubation with specific primary antibodies for 1 h. The cells were then incubated for 45 min with AlexaFluor-conjugated secondary antibodies, or with AlexaFluor568 phalloidin to visualize actin filaments. Nuclei were counterstained with Hoechst 34580 (H21486; ThermoFisher Scientific). Coverslips were mounted onto microscopy slides (Globe Scientific INC, Mahwah, NJ, USA) in Prolong Glass Antifade mounting medium (ThermoFisher Scientific). Images were acquired on an LSM Olympus FV1000 spectral SIM confocal microscope (Tokyo, JP) using the Olympus FluoView FV-10 ASW software v4.2. Image analyses were performed on CellProfiler v4.0.7 [31] using Mander’s overlap to quantify colocalization of MET with the different subcellular compartment markers.

### Flow cytometry

The surface expression of MET was determined by flow cytometry. Cells were plated in 35-mm dish and harvested after 48 h using cold flow cytometry buffer (PBS 1X; 2% FBS) and stained using AlexaFluor488-coupled anti-MET antibody diluted in flow cytometry buffer. Validated DsiRNAs (IDT, Coralville, IA, USA) were used to down-regulate MET expression in HCT116 cells, which were included as a control. Data were acquired using a Guava EasyCyte Mini cytometer (EMD Millipore) and analyzed using Kaluza Analysis v2.2 (Beckman Coulter, Brea, CA, USA).

### EdU incorporation assay

Serum-starved cells were incubated with or without HGF (50 ng/mL) for 12 h, before the addition of EdU (15µM, 5-ethynyl-2’-deoxyuridine) to the medium for 90 min. Then, cells were harvested by trypsinization and fixed in 4% formaldehyde. The EdU incorporation was determined with the Click-iT EdU Alexa Fluor 647 Flow Cytometry Assay kit (C10424; Thermo Fisher Scientific) following the manufacturer’s instructions. The DNA content was assessed by DAPI (4′,6-diamidino-2-phenylindole; Sigma-Aldrich) staining. Cells were analyzed by an automatic 96-well-plate loader-equipped CytoFLEX flow cytometer (Beckman Coulter).

### Cell growth, scatter, and migration/wound healing assays

The methodology for cell growth was previously described [32]. Scattering assay consisted of stimulating serum-starved cells with HGF (50 ng/mL) for 24 h, documenting the cells’ response before and after stimulation by phase-contrast imaging. For wound healing assays, cells were grown until confluence was reached in a 100 mm plate. Then, a scratch was made by scraping the cell monolayer with a 10 µL micropipette tip and the plate was washed with DMEM to eliminate floating cells. Cultures were maintained in serum-free DMEM in the presence or not of HGF (100 ng/mL) and photographs were taken at 0 and 24 h. The migration distance, representing the percentage of wound healing, was calculated from the scratch area measurements using ImageJ [33].

### Statistical analyses

Results were analyzed using the GraphPad Prism v.9.0 software, comparing parental and *SNX1/2* KO cells using the *t*-test. Microscopy images were obtained from three different fields containing approximately 60 cells from three independent experiments and the colocalization was analyzed using a two-way ANOVA with Sidak correction. The significance was expressed by the *p-*value (*, *p* ≤ 0.05; **, *p* ≤ 0.01; ***, *p* ≤ 0.001; ****, *p* ≤ 0.001).

## Supporting information

Manuscript Supplementals

## DATA AVAILABILITY

All data are included in the manuscript and/or supplemental file.

## ACKNOWLEDGEMENTS

This work was supported by the Canadian Institutes for Health Research (CIHR) PJT-162354 (to J-BD, CS, and CLL). LGD was supported by an IRCUS excellence studentship and AD by a CIHR studentship award. J-BD, CS and CLL are members of the FRQS-funded Centre de Recherche du CHUS.

## AUTHORS CONTRIBUTION

**LGD** designed the experiments, performed the research, analyzed the data, and wrote the manuscript.

**AD** designed the experiments, performed the research, analyzed the data.

**SL** performed the research.

**MGD** performed the research.

**GB** performed the research.

**CLL** designed the experiments, analyzed the data, and wrote the manuscript.

**CS** designed the experiments, analyzed the data, and wrote the manuscript.

**J-BD** designed the experiments, analyzed the data, and wrote the manuscript.

## References

[1] Barrow-McGee R, Kermorgant S. Met endosomal signalling: in the right place, at the right time. Int J Biochem Cell Biol 2014;49:69–74. 10.1016/j.biocel.2014.01.009.

[2] Bergeron JJM, Di Guglielmo GM, Dahan S, Dominguez M, Posner BI. Spatial and Temporal Regulation of Receptor Tyrosine Kinase Activation and Intracellular Signal Transduction. Annu Rev Biochem 2016;85:573–97. 10.1146/annurev-biochem-060815-014659.

[3] Crilly SE, Puthenveedu MA. Compartmentalized GPCR Signaling from Intracellular Membranes. J Membr Biol 2021;254:259–71. 10.1007/s00232-020-00158-7.

[4] Vieira AV, Lamaze C, Schmid SL. Control of EGF Receptor Signaling by Clathrin-Mediated Endocytosis. Science 1996;274:2086–9. 10.1126/science.274.5295.2086.

[5] Gormal RS, Martinez-Marmol R, Brooks AJ, Meunier FA. Location, location, location: Protein kinase nanoclustering for optimised signalling output. eLife 2024;13:e93902. 10.7554/eLife.93902.

[6] Murphy JE, Padilla BE, Hasdemir B, Cottrell GS, Bunnett NW. Endosomes: A legitimate platform for the signaling train. Proc Natl Acad Sci 2009;106:17615–22. 10.1073/pnas.0906541106.

[7] Daaka Y, Luttrell LM, Ahn S, Rocca GJD, Ferguson SSG, Caron MG, et al. Essential Role for G Protein-coupled Receptor Endocytosis in the Activation of Mitogen-activated Protein Kinase *. J Biol Chem 1998;273:685–8. 10.1074/jbc.273.2.685.

[8] Pavlos NJ, Friedman PA. GPCR Signaling and Trafficking: The Long and Short of It. Trends Endocrinol Metab 2017;28:213–26. 10.1016/j.tem.2016.10.007.

[9] Sung H, Ferlay J, Siegel RL, Laversanne M, Soerjomataram I, Jemal A, et al. Global Cancer Statistics 2020: GLOBOCAN Estimates of Incidence and Mortality Worldwide for 36 Cancers in 185 Countries. CA Cancer J Clin 2021;71:209–49. 10.3322/caac.21660.

[10] Comoglio PM, Trusolino L, Boccaccio C. Known and novel roles of the MET oncogene in cancer: a coherent approach to targeted therapy. Nat Rev Cancer 2018;18:341–58. 10.1038/s41568-018-0002-y.

[11] Saucier C, Rivard N. Epithelial Cell Signalling in Colorectal Cancer Metastasis. In: Beauchemin N, Huot J, editors. Metastasis Colorectal Cancer, Dordrecht: Springer Netherlands; 2010, p. 205–41. 10.1007/978-90-481-8833-8_8.

[12] Trusolino L, Bertotti A, Comoglio PM. MET signalling: principles and functions in development, organ regeneration and cancer. Nat Rev Mol Cell Biol 2010;11:834–48. 10.1038/nrm3012.

[13] Boccaccio C, Andò M, Tamagnone L, Bardelli A, Michieli P, Battistini C, et al. Induction of epithelial tubules by growth factor HGF depends on the STAT pathway. Nature 1998;391:285–8. 10.1038/34657.

[14] Kermorgant S, Zicha D, Parker PJ. PKC controls HGF-dependent c-Met traffic, signalling and cell migration. EMBO J 2004;23:3721–34. 10.1038/sj.emboj.7600396.

[15] Abella JV, Peschard P, Naujokas MA, Lin T, Saucier C, Urbé S, et al. Met/Hepatocyte Growth Factor Receptor Ubiquitination Suppresses Transformation and Is Required for Hrs Phosphorylation. Mol Cell Biol 2005;25:9632–45. 10.1128/MCB.25.21.9632-9645.2005.

[16] Parachoniak CA, Luo Y, Abella JV, Keen JH, Park M. GGA3 Functions as a Switch to Promote Met Receptor Recycling, Essential for Sustained ERK and Cell Migration. Dev Cell 2011;20:751–63. 10.1016/j.devcel.2011.05.007.

[17] Kermorgant S, Parker PJ. Receptor trafficking controls weak signal delivery: a strategy used by c-Met for STAT3 nuclear accumulation. J Cell Biol 2008;182:855–63. 10.1083/jcb.200806076.

[18] Joffre C, Barrow R, Ménard L, Calleja V, Hart IR, Kermorgant S. A direct role for Met endocytosis in tumorigenesis. Nat Cell Biol 2011;13:827–37. 10.1038/ncb2257.

[19] Tan Y, You H, Wu C, Altomare DA, Testa JR. Appl1 Is Dispensable for Mouse Development, and Loss of Appl1 Has Growth Factor-selective Effects on Akt Signaling in Murine Embryonic Fibroblasts *. J Biol Chem 2010;285:6377–89. 10.1074/jbc.M109.068452.

[20] Teasdale RD, Collins BM. Insights into the PX (phox-homology) domain and SNX (sorting nexin) protein families: structures, functions and roles in disease. Biochem J 2012;441:39–59. 10.1042/BJ20111226.

[21] Wassmer T, Attar N, Bujny MV, Oakley J, Traer CJ, Cullen PJ. A loss-of-function screen reveals SNX5 and SNX6 as potential components of the mammalian retromer. J Cell Sci 2007;120:45–54. 10.1242/jcs.03302.

[22] Haft CR, de la Luz Sierra M, Barr VA, Haft DH, Taylor SI. Identification of a family of sorting nexin molecules and characterization of their association with receptors. Mol Cell Biol 1998;18:7278–87. 10.1128/mcb.18.12.7278.

[23] Kvainickas A, Jimenez-Orgaz A, Nägele H, Hu Z, Dengjel J, Steinberg F. Cargo-selective SNX-BAR proteins mediate retromer trimer independent retrograde transport. J Cell Biol 2017;216:3677–93. 10.1083/jcb.201702137.

[24] Yang Z, Feng Z, Li Z, Teasdale RD. Multifaceted Roles of Retromer in EGFR Trafficking and Signaling Activation. Cells 2022;11:3358. 10.3390/cells11213358.

[25] Schaaf CP, Benzing J, Schmitt T, Erz DHR, Tewes M, Bartram CR, et al. Novel interaction partners of the TPR/MET tyrosine kinase. FASEB J Off Publ Fed Am Soc Exp Biol 2005;19:267–9. 10.1096/fj.04-1558fje.

[26] Ogi S, Fujita H, Kashihara M, Yamamoto C, Sonoda K, Okamoto I, et al. Sorting nexin 2-mediated membrane trafficking of c-Met contributes to sensitivity of molecular-targeted drugs. Cancer Sci 2013;104:573–83. 10.1111/cas.12117.

[27] Duclos CM, Champagne A, Carrier JC, Saucier C, Lavoie CL, Denault J-B. Caspase-mediated proteolysis of the sorting nexin 2 disrupts retromer assembly and potentiates Met/hepatocyte growth factor receptor signaling. Cell Death Discov 2017;3:16100. 10.1038/cddiscovery.2016.100.

[28] Bian Z, Feng Y, Xue Y, Hu Y, Wang Q, Zhou L, et al. Down-regulation of SNX1 predicts poor prognosis and contributes to drug resistance in colorectal cancer. Tumour Biol 2016;37:6619–25. 10.1007/s13277-015-3814-3.

[29] Nishimura Y, Takiguchi S, Ito S, Itoh K. Evidence that depletion of the sorting nexin 1 by siRNA promotes HGF-induced MET endocytosis and MET phosphorylation in a gefitinib-resistant human lung cancer cell line. Int J Oncol 2014;44:412–26. 10.3892/ijo.2013.2194.

[30] Bury AF. Analysis of protein and peptide mixtures: Evaluation of three sodium dodecyl sulphate-polyacrylamide gel electrophoresis buffer systems. J Chromatogr A 1981;213:491–500. 10.1016/S0021-9673(00)80500-2.

[31] Stirling DR, Swain-Bowden MJ, Lucas AM, Carpenter AE, Cimini BA, Goodman A. CellProfiler 4: improvements in speed, utility and usability. BMC Bioinformatics 2021;22:433. 10.1186/s12859-021-04344-9.

[32] McManus S, Chababi W, Arsenault D, Dubois CM, Saucier C. Dissecting Oncogenic RTK Pathways in Colorectal Cancer Initiation and Progression. In: Beaulieu J-F, editor. Colorectal Cancer Methods Protoc., New York, NY: Springer New York; 2018, p. 27–42. 10.1007/978-1-4939-7765-9_2.

[33] Schneider CA, Rasband WS, Eliceiri KW. NIH Image to ImageJ: 25 years of image analysis. Nat Methods 2012;9:671–5. 10.1038/nmeth.2089.

[34] Gullapalli A, Garrett TA, Paing MM, Griffin CT, Yang Y, Trejo J. A Role for Sorting Nexin 2 in Epidermal Growth Factor Receptor Down-regulation: Evidence for Distinct Functions of Sorting Nexin 1 and 2 in Protein Trafficking. Mol Biol Cell 2004;15:2143–55. 10.1091/mbc.e03-09-0711.

[35] Mu F-T, Callaghan JM, Steele-Mortimer O, Stenmark H, Parton RG, Campbell PL, et al. EEA1, an Early Endosome-Associated Protein.: EEA1 IS A CONSERVED α-HELICAL PERIPHERAL MEMBRANE PROTEIN FLANKED BY CYSTEINE “FINGERS” AND CONTAINS A CALMODULIN-BINDING IQ MOTIF ∗. J Biol Chem 1995;270:13503–11. 10.1074/jbc.270.22.13503.

[36] Meresse S, Gorvel JP, Chavrier P. The rab7 GTPase resides on a vesicular compartment connected to lysosomes. J Cell Sci 1995;108:3349–58. 10.1242/jcs.108.11.3349.

[37] Fukuda M. Lysosomal membrane glycoproteins. Structure, biosynthesis, and intracellular trafficking. J Biol Chem 1991;266:21327–30. 10.1016/S0021-9258(18)54636-6.

[38] Kermorgant S, Zicha D, Parker PJ. Protein Kinase C Controls Microtubule-based Traffic but Not Proteasomal Degradation of c-Met *. J Biol Chem 2003;278:28921–9. 10.1074/jbc.M302116200.

[39] Bali PK, Zak O, Aisen P. A new role for the transferrin receptor in the release of iron from transferrin. Biochemistry 1991;30:324–8. 10.1021/bi00216a003.

[40] Ahmed D, Eide PW, Eilertsen IA, Danielsen SA, Eknæs M, Hektoen M, et al. Epigenetic and genetic features of 24 colon cancer cell lines. Oncogenesis 2013;2:e71. 10.1038/oncsis.2013.35.

[41] Furge KA, Zhang Y-W, Vande Woude GF. Met receptor tyrosine kinase: enhanced signaling through adapter proteins. Oncogene 2000;19:5582–9. 10.1038/sj.onc.1203859.

[42] Kaufmann SH, Desnoyers S, Ottaviano Y, Davidson NE, Poirier GG. Specific proteolytic cleavage of poly(ADP-ribose) polymerase: an early marker of chemotherapy-induced apoptosis. Cancer Res 1993;53:3976–85.

[43] Kurten RC, Cadena DL, Gill GN. Enhanced degradation of EGF receptors by a sorting nexin, SNX1. Science 1996;272:1008–10. 10.1126/science.272.5264.1008.

[44] Simonetti B, Danson CM, Heesom KJ, Cullen PJ. Sequence-dependent cargo recognition by SNX-BARs mediates retromer-independent transport of CI-MPR. J Cell Biol 2017;216:3695–712. 10.1083/jcb.201703015.

[45] Griffin CT, Trejo J, Magnuson T. Genetic evidence for a mammalian retromer complex containing sorting nexins 1 and 2. Proc Natl Acad Sci 2005;102:15173–7. 10.1073/pnas.0409558102.

[46] Mathieson T, Franken H, Kosinski J, Kurzawa N, Zinn N, Sweetman G, et al. Systematic analysis of protein turnover in primary cells. Nat Commun 2018;9:689. 10.1038/s41467-018-03106-1.

[47] Pomerleau V, Landry M, Bernier J, Vachon PH, Saucier C. Met receptor-induced Grb2 or Shc signals both promote transformation of intestinal epithelial cells, albeit they are required for distinct oncogenic functions. BMC Cancer 2014;14:240. 10.1186/1471-2407-14-240.

[48] Zhan X-Y, Zhang Y, Zhai E, Zhu Q-Y, He Y. Sorting nexin-1 is a candidate tumor suppressor and potential prognostic marker in gastric cancer. PeerJ 2018;6:e4829. 10.7717/peerj.4829.

[49] Zhou Q, Li J, Ge C, Chen J, Tian W, Tian H. SNX5 suppresses clear cell renal cell carcinoma progression by inducing CD44 internalization and epithelial-to-mesenchymal transition. Mol Ther - Oncolytics 2022;24:87–100. 10.1016/j.omto.2021.12.002.

[50] Grant BD, Donaldson JG. Pathways and mechanisms of endocytic recycling. Nat Rev Mol Cell Biol 2009;10:597–608. 10.1038/nrm2755.

[51] Kalaidzidis I, Miaczynska M, Brewińska-Olchowik M, Hupalowska A, Ferguson C, Parton RG, et al. APPL endosomes are not obligatory endocytic intermediates but act as stable cargo-sorting compartments. J Cell Biol 2015;211:123–44. 10.1083/jcb.201311117.

[52] Jean-Alphonse F, Bowersox S, Chen S, Beard G, Puthenveedu MA, Hanyaloglu AC. Spatially restricted G protein-coupled receptor activity via divergent endocytic compartments. J Biol Chem 2014;289:3960–77. 10.1074/jbc.M113.526350.

[53] Lee J-R, Hahn H-S, Kim Y-H, Nguyen H-H, Yang J-M, Kang J-S, et al. Adaptor protein containing PH domain, PTB domain and leucine zipper (APPL1) regulates the protein level of EGFR by modulating its trafficking. Biochem Biophys Res Commun 2011;415:206–11. 10.1016/j.bbrc.2011.10.064.

[54] York HM, Patil A, Moorthi UK, Kaur A, Bhowmik A, Hyde GJ, et al. Rapid whole cell imaging reveals a calcium-APPL1-dynein nexus that regulates cohort trafficking of stimulated EGF receptors. Commun Biol 2021;4:1–13. 10.1038/s42003-021-01740-y.

[55] Li J, Mao X, Dong LQ, Liu F, Tong L. Crystal Structures of the BAR-PH and PTB Domains of Human APPL1. Structure 2007;15:525–33. 10.1016/j.str.2007.03.011.

[56] Franke TF, Tartof KD, Tsichlis PN. The SH2-like Akt homology (AH) domain of c-akt is present in multiple copies in the genome of vertebrate and invertebrate eucaryotes. Cloning and characterization of the Drosophila melanogaster c-akt homolog Dakt1. Oncogene 1994;9:141–8.

[57] Kavanaugh WM, Williams LT. An Alternative to SH2 Domains for Binding Tyrosine-Phosphorylated Proteins. Science 1994;266:1862–5. 10.1126/science.7527937.

[58] Zoncu R, Perera RM, Balkin DM, Pirruccello M, Toomre D, De Camilli P. A Phosphoinositide Switch Controls the Maturation and Signaling Properties of APPL Endosomes. Cell 2009;136:1110–21. 10.1016/j.cell.2009.01.032.

[59] York HM, Joshi K, Wright CS, Kreplin LZ, Rodgers SJ, Moorthi UK, et al. Deterministic early endosomal maturations emerge from a stochastic trigger-and-convert mechanism. Nat Commun 2023;14:4652. 10.1038/s41467-023-40428-1.

[60] Schenck A, Goto-Silva L, Collinet C, Rhinn M, Giner A, Habermann B, et al. The Endosomal Protein Appl1 Mediates Akt Substrate Specificity and Cell Survival in Vertebrate Development. Cell 2008;133:486–97. 10.1016/j.cell.2008.02.044.

[61] Deepa SS, Dong LQ. APPL1: role in adiponectin signaling and beyond. Am J Physiol Endocrinol Metab 2009;296:E22–36. 10.1152/ajpendo.90731.2008.

[62] Zhou J, Liu H, Zhou S, He P, Liu X. Adaptor protein APPL1 interacts with EGFR to orchestrate EGF-stimulated signaling. Sci Bull 2016;61:1504–12. 10.1007/s11434-016-1157-0.

[63] Sondka Z, Dhir NB, Carvalho-Silva D, Jupe S, Madhumita, McLaren K, et al. COSMIC: a curated database of somatic variants and clinical data for cancer. Nucleic Acids Res 2024;52:D1210–7. 10.1093/nar/gkad986.

[64] Hanahan D, Weinberg RA. Hallmarks of cancer: the next generation. Cell 2011;144:646–74. 10.1016/j.cell.2011.02.013.

[65] Chandarlapaty S. Negative Feedback and Adaptive Resistance to the Targeted Therapy of Cancer. Cancer Discov 2012;2:311–9. 10.1158/2159-8290.CD-12-0018.

[66] Royal I, Park M. Hepatocyte Growth Factor-induced Scatter of Madin-Darby Canine Kidney Cells Requires Phosphatidylinositol 3-Kinase (*). J Biol Chem 1995;270:27780–7. 10.1074/jbc.270.46.27780.

[67] Ridley AJ, Comoglio PM, Hall A. Regulation of scatter factor/hepatocyte growth factor responses by Ras, Rac, and Rho in MDCK cells. Mol Cell Biol 1995;15:1110–22.

[68] Tewari M, Quan LT, O’Rourke K, Desnoyers S, Zeng Z, Beidler DR, et al. Yama/CPP32 beta, a mammalian homolog of CED-3, is a CrmA-inhibitable protease that cleaves the death substrate poly(ADP-ribose) polymerase. Cell 1995;81:801–9. 10.1016/0092-8674(95)90541-3.

[69] Germain M, Affar EB, D’Amours D, Dixit VM, Salvesen GS, Poirier GG. Cleavage of Automodified Poly(ADP-ribose) Polymerase during Apoptosis: EVIDENCE FOR INVOLVEMENT OF CASPASE-7 *. J Biol Chem 1999;274:28379–84. 10.1074/jbc.274.40.28379.

[70] Cardone MH, Roy N, Stennicke HR, Salvesen GS, Franke TF, Stanbridge E, et al. Regulation of cell death protease caspase-9 by phosphorylation. Science 1998;282:1318–21. 10.1126/science.282.5392.1318.

[71] Allan LA, Morrice N, Brady S, Magee G, Pathak S, Clarke PR. Inhibition of caspase-9 through phosphorylation at Thr 125 by ERK MAPK. Nat Cell Biol 2003;5:647–54. 10.1038/ncb1005.

[72] Ozören N, El-Deiry WS. Defining characteristics of Types I and II apoptotic cells in response to TRAIL. Neoplasia N Y N 2002;4:551–7. 10.1038/sj.neo.7900270.

[73] Atwell B, Chen C-Y, Christofferson M, Montfort WR, Schroeder J. Sorting nexin-dependent therapeutic targeting of oncogenic epidermal growth factor receptor. Cancer Gene Ther 2023;30:267–76. 10.1038/s41417-022-00541-7.

