## Supplementary material for "Spatiotemporal regulation of the hepatocyte growth factor receptor MET activity by sorting nexins 1/2 in HCT116 colorectal cancer cells": Manuscript Supplementals

**A**

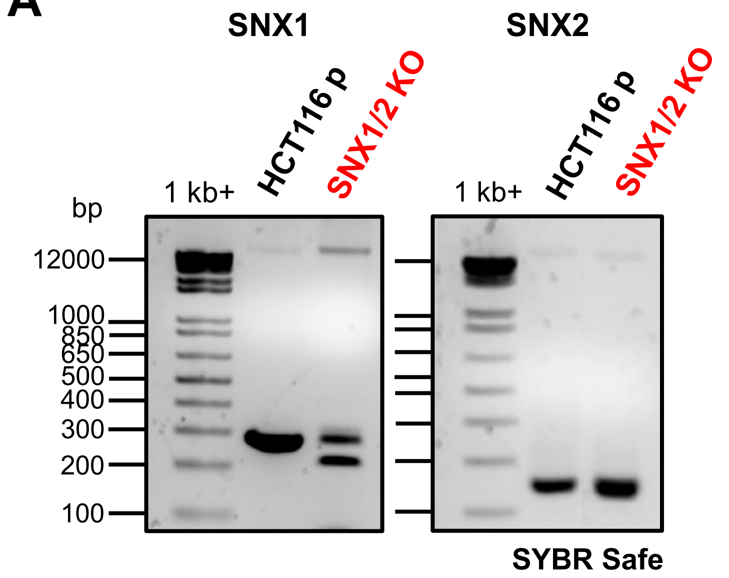


**B**

| Cells | SNX1 sequence | SNX2 sequence |
|---|---|---|
| HCT116 p | wt | wt |
| SNX1/2 KO | 68 nt deletion<br>11 nt deletion | 4 nt deletion<br>1 nt insertion |

**Suppl. Figure 1: The knockout of *SNX1* or *SNX2* in HCT116 cells was validated by PCR and sequencing. A** Genomic DNA from parental and *SNX1/2* KO cells were subjected to PCR amplification of *SNX1* or *SNX2* specific sequences (including the CRISPR/Cas9 target region), using the primer pairs SNX1-F: 5'-GCAGTGTCAGCTGATTTGTCC-3'; SNX1-R: 5'-AACAGAAGCTTACGCGGACT-3'; SNX2-F: 5'-TGATGCTAAATTGTGCATTGCC-3'; SNX2-R: 5'-TGCAGCAAAATGTGACCATGT-3', and the PCR products were next sequenced. **B** The table shows a summary of the modifications in the sequence of *SNX1* and *SNX2* resulting in the depletion of both proteins.

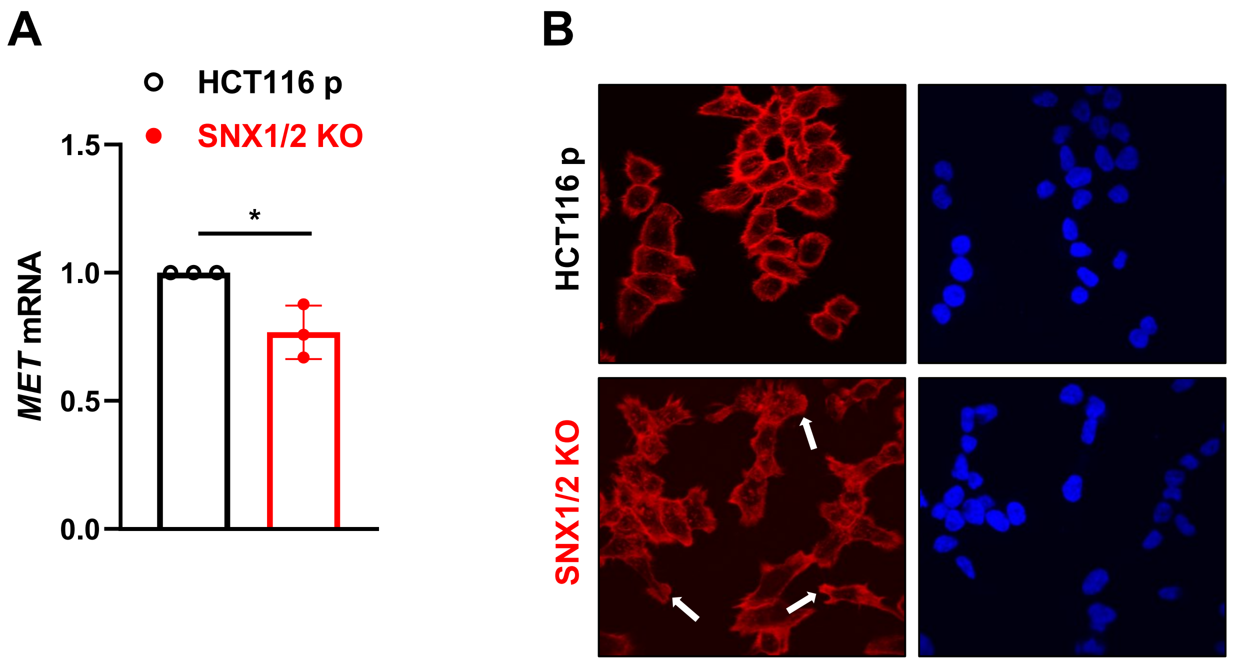


**Suppl. Figure 2: The knockout of SNX1 and SNX2 decreases *MET* mRNA levels and induces actin reorganization in HCT116 cells. A** The level of *MET* mRNA was determined by qRT-PCR using the primers 5'-TGGCTACACACTGGTTATCACTGG-3' and 5'-ACTGGAAATGTCTGCAGCCCAA-3'. MRPL19 (mitochondrial ribosomal protein L19), PUM1 (pumilio RNA-binding family member 1), and YWHAZ (tyrosine 3-monooxygenase/tryptophan 5-monooxygenase activation protein zeta) were used as reference genes. The bar graph represents the mean ± SD of three independent experiments, normalized to the parental cells (N=3). **B** Confocal microscopy images for polymerized actin staining (phalloidin). Nuclei were counterstained with Hoechst. Arrows indicate membrane protrusions. Scale 10 µm; 60X objective.

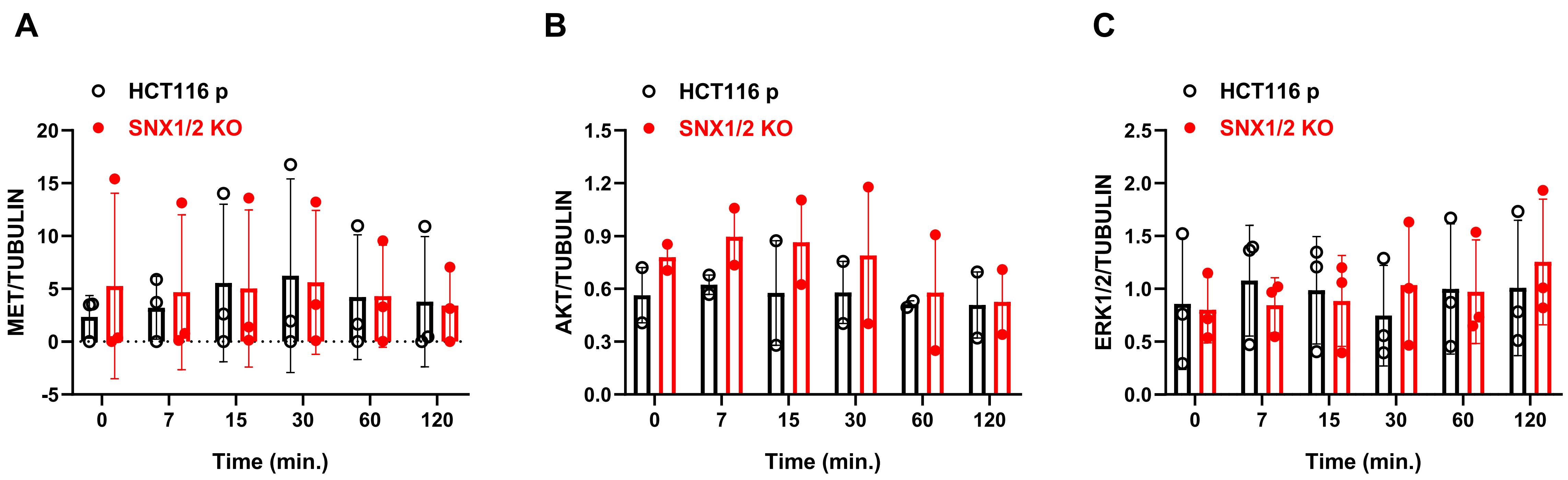


**Suppl. Figure 3: The levels of total MET, ERK1/2, and AKT do not differ between parental and *SNX1/2* KO cells upon HGF stimulation.** Serum-starved cells were stimulated with 50 ng/mL HGF for the indicated period. Graphs show the densitometric quantification of data from **Figure 5A**, expressed as the ratio of total **A** MET, **B** ERK1/2, and **C** AKT proteins, normalized to the loading control tubulin. Data correspond to the mean ± SD (N=3).

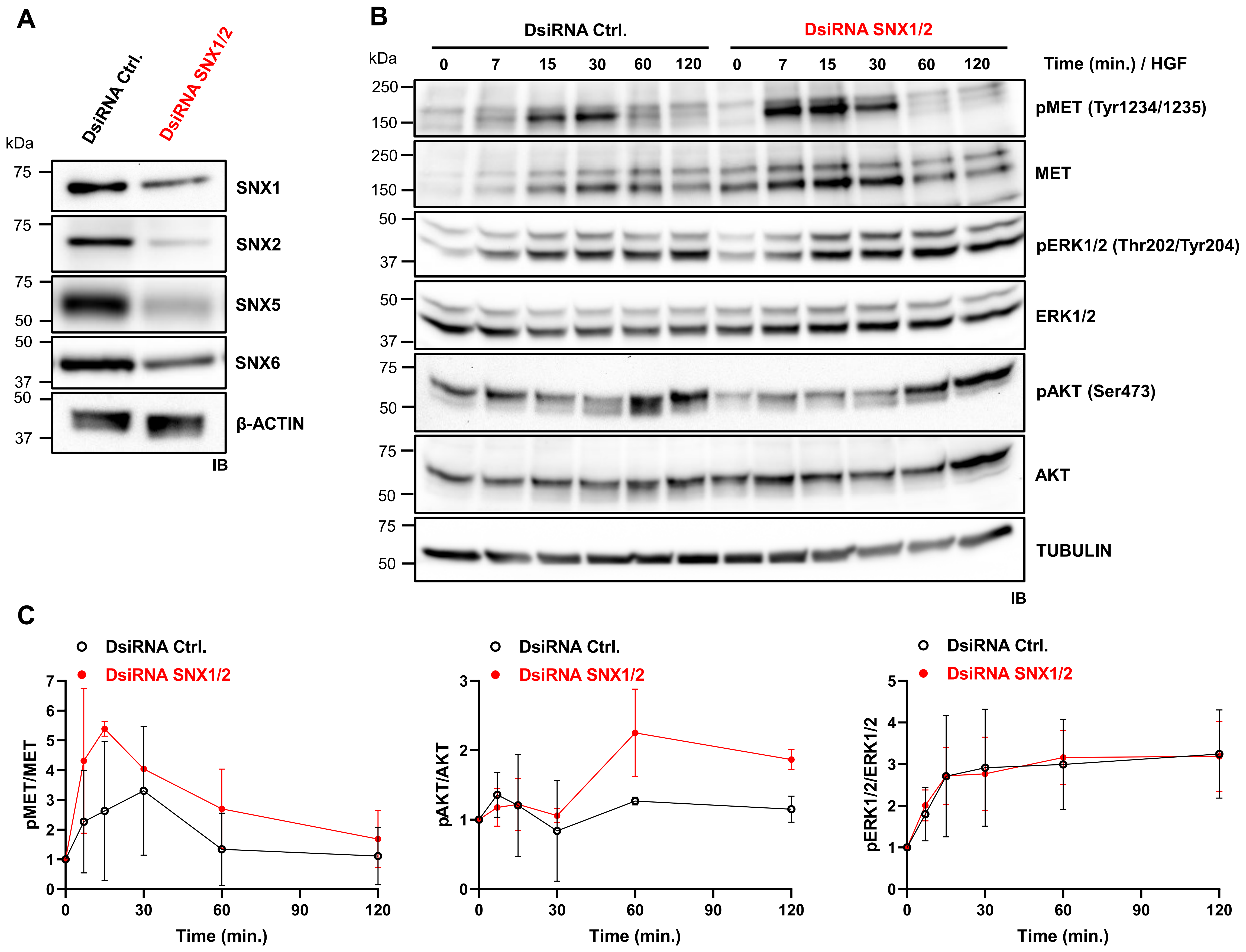


**Suppl. Figure 4: The *SNX1* and *SNX2* knockdown potentiates the phosphorylation of MET and AKT upon HGF stimulation. A** Validation of reduced SNX1 and SNX2 protein expression following DsiRNAs transfection, as well as of the resulting down-regulation of SNX5 and SNX6, by immunoblotting. **B** DsiRNA-transfected cells were serum-starved and stimulated with HGF (50 ng/mL) for the indicated period. Phosphorylation of MET and the downstream effectors AKT and ERK1/2 was analyzed by immunoblotting. Tubulin was used as a loading control. **C** Graphs show the densitometric quantification of data from **B** expressed as the ratio of phosphorylated protein/total protein and normalized to the unstimulated sample. Data corresponds to the mean ± SD (N=3).

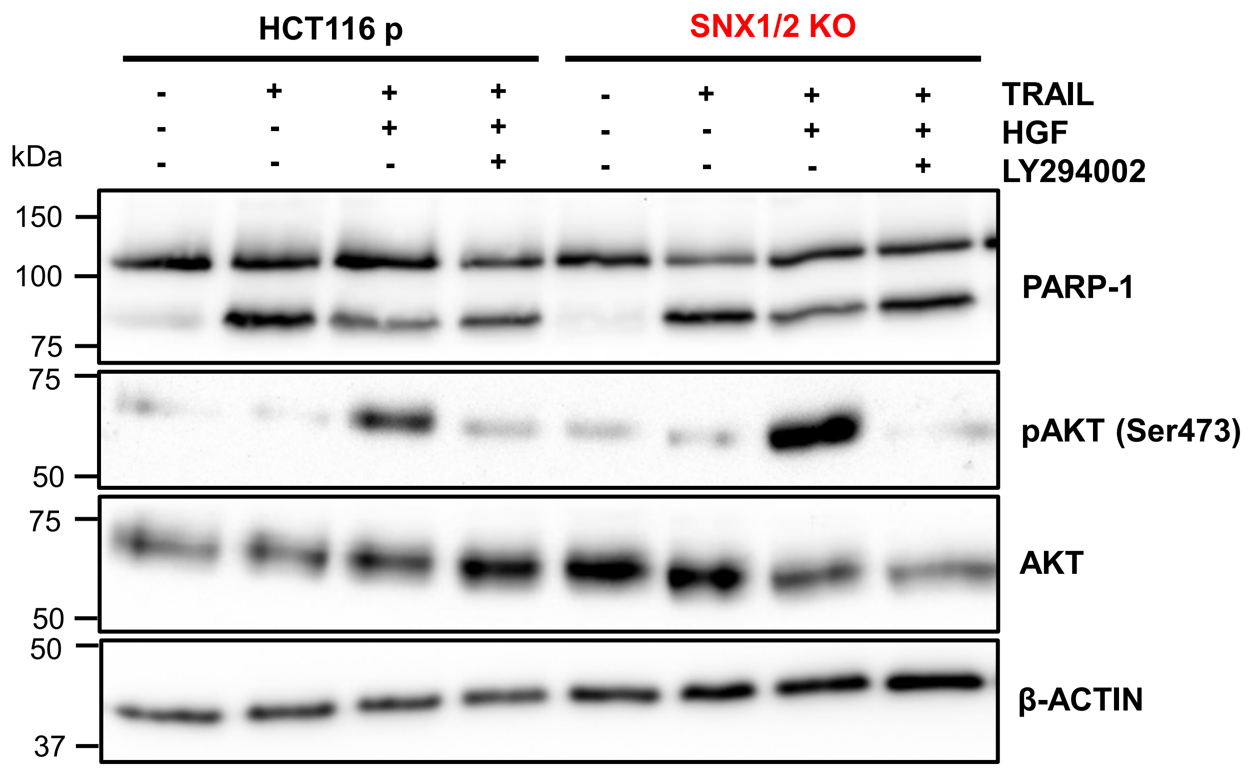


**Suppl. Figure 5: The protective effect from TRAIL-induced apoptosis depends on MET-PI3K-AKT activation.** Apoptosis was induced by treating cells with TRAIL (50 ng/mL). Simultaneously, cells were treated with HGF (100 ng/mL) alone or with the PI3K inhibitor LY294002 (10 μM). The cleavage of PARP-1 was determined by immunoblotting; actin was used as a loading control.

**Table S1. Antibodies used.**

| Antibodies for immunoblotting | | | |
|---|---|---|---|
| **Antibody** | **Dilution** | **Catalog number** | **Source** |
| MET (D-4) | 1:1000 | sc-514148 | Santa Cruz Biotechnology (Dallas, TX, USA) |
| SNX5 (F-11) | 1:1000 | sc-515215 | Santa Cruz Biotechnology |
| SNX6 (D-5) | 1:1000 | sc-365965 | Santa Cruz Biotechnology |
| SNX2 | 1:1000 | Y00322-002 | Immune Biosolutions (Sherbrooke, QC, Canada) |
| SNX1 (51) | 0.25 µg/mL | 611482 | BD Biosciences (Franklin Lakes, NJ, USA) |
| PARP-1 (C2-10) | 556362 | 1:7500 | BD Biosciences |
| Phospho-MET (Tyr1234/1235) (D26) | 1:1000 | 3077 | Cell Signaling Biotechnology (Danvers, MA, USA) |
| Phospho-p44/42 MAPK (Erk1/2) (Thr202/Tyr204) | 1:2000 | 4370 | Cell Signaling Technology |
| p44/42 MAPK (Erk1/2) | 1:1000 | 4695 | Cell Signaling Technology |
| Phospho-AKT (Ser473) | 1:500 | 9271 | Cell Signaling Technology |
| AKT | 1:500 | 9272 | Cell Signaling Technology |
| E-cadherin (36) | 0.25 µg/mL | 610181 | BD Biosciences |
| Actin (AC-40) | 1:5000 | A3853 | Sigma-Aldrich (St-Louis, MO, USA) |
| α-tubulin (B-5-1-2) | 1:5000 | T5168 | Sigma-Aldrich |
| Anti-mouse IgG, HRP-linked | 1:5000 | 7076 | Cell Signaling Technology |
| Anti-rabbit IgG, HRP-linked | 1:5000 | 7074 | Cell Signaling Technology |
| Anti-chicken IgG, HRP-linked | 1:3000 | Y00008-002 | Immune Biosolutions |

| Antibodies for immunofluorescence | | | |
|---|---|---|---|
| **Antibody** | **Dilution** | **Catalog number** | **Source** |
| MET (L6E7) | 1:1000 | 8741 | Cell Signaling Technology |

| | | | |
|---|---|---|---|
| EEA1 | 10 µg/mL | PA1-063A | Thermo Fisher Scientific (Waltham, MA, USA) |
| CD71 (D7G9X) | 1:100 | 13113 | Cell Signaling Technology |
| LAMP1 (D2D11) | 5 µg/mL | 9091 | Cell Signaling Technology |
| RAB7 (D95F2) | 1:50 | 9367 | Cell Signaling Technology |
| GGA3 | 1 µg/mL | PA5-82888 | Thermo Fisher Scientific |
| Goat anti-Rabbit IgG (H+L) Cross-Adsorbed Secondary Antibody, Alexa Fluor™ 405 | 12 ng/mL | A31556 | Thermo Fisher Scientific |
| Donkey anti-Rabbit IgG (H+L) Highly Cross-Adsorbed Secondary Antibody, Alexa Fluor™ 488 | 12 ng/mL | A21206 | Thermo Fisher Scientific |
| Donkey anti-Rabbit IgG (H+L) Highly Cross-Adsorbed Secondary Antibody, Alexa Fluor™ 594 | 12 ng/mL | A21207 | Thermo Fisher Scientific |
| Donkey anti-Mouse IgG (H+L) Highly Cross-Adsorbed Secondary Antibody, Alexa Fluor™ 488 | 12 ng/mL | A21202 | Thermo Fisher Scientific |
| Donkey anti-Mouse IgG (H+L) Highly Cross-Adsorbed Secondary Antibody, Alexa Fluor™ 594 | 12 ng/mL | A21203 | Thermo Fisher Scientific |
| Alexa Fluor™ 568 Phalloidin | 1:1000 | A12380 | Thermo Fisher Scientific |

| Antibodies for cytometry | | | |
|---|---|---|---|
| **Antibody** | **Dilution** | **Catalog number** | **Source** |
| Human HGFR/c-MET Alexa Fluor® 488-conjugated | 5 µL/$10^6$ cells | FAB3582G | R&D systems (Minneapolis, MN, USA) |
